# Fate mapping quantifies the dynamics of B cell development and activation throughout life

**DOI:** 10.1101/871624

**Authors:** Melissa Verheijen, Sanket Rane, Claire Pearson, Andrew J. Yates, Benedict Seddon

## Abstract

Follicular mature (FM) and germinal centre (GC) B cells underpin humoral immunity but the dynamics of their generation and maintenance are not clearly defined. Here we exploited a fate-mapping system in mice that tracks B cells as they develop into peripheral subsets, together with a cell division fate reporter mouse and mathematical models. We find that FM cells are kinetically homogeneous, recirculate freely, continually replenished from transitional populations, and self-renew rarely. In contrast, GC B cell lineages persist for weeks with rapid turnover and site-specific dynamics. Those in the spleen derive from transitional cells and are kinetically homogeneous, while those in lymph nodes derive from FM B cells and comprise both transient and persistent clones. These differences likely derive from the nature of antigen exposure at the different sites. Our integrative approach also reveals how the host environment drives cell-extrinsic, age-related changes in B cell homeostasis.

## Introduction

The ability to mount effective humoral immune responses throughout life is critical for normal antibody-mediated protection and healthy aging (Gibson et al., 2009, Frasca et al., 2011). B cells are generated in the bone marrow and enter the spleen where they complete development as transitional cells, characterised by induction of CD23 and IgD, together with downregulation of IgM and AA4.1 (Allman et al., 2001, Loder et al., 1999). These markers identify three stages of transitional cell maturation. During the T1 stage, IgM^hi^ CD23^low^ B cells with autoreactive BCRs undergo negative selection (Petro et al., 2002, Su and Rawlings, 2002). During the T2 stage, CD23^hi^IgD^hi^ cells commit to either a follicular B cell fate, progressing through the IgM^low^ T3 stage, or are diverted to develop into marginal zone B cells, losing expression of CD23, upregulating IgM and expressing CD21 (Petro et al., 2002, Su and Rawlings, 2002, Lam et al., 1997, Cariappa, 2009, Torres et al., 1996). Follicular mature (FM) B cells recirculate between lymph nodes and spleen, where cognate encounter with antigen triggers activation and the development of germinal centre reactions. In deliberately challenged mice, antigen-specific germinal centre (GC) B cells divide extensively and undergo affinity maturation (Mesin et al., 2016, Basso and Dalla-Favera, 2015, De Silva and Klein, 2015). However, GC B cells are present throughout a mouse’s lifetime even in the absence of deliberate immunological challenge. The origin and dynamics of these constitutive GC reactions are not well characterised.

While the establishment of peripheral B cell subsets relies upon *de novo* generation in the bone marrow, it is unclear to what degrees the processes of influx of new cells, proliferative renewal, and cell loss (turnover) combine to maintain B cell subsets at or close to dynamic equilibrium, and how these process may change throughout life. Much of our insights into these processes derive from DNA labelling experiments using BrdU. Numbers of immature B cells in the spleen decline with age, and it has been inferred from BrdU labelling that this decline derives from a loss of efficiency of pre-transitional B cell development, rather than any decrease in the rate of production of B cell progenitors in the bone marrow (Kline et al., 1999, Shahaf et al., 2006). In adult mice, it has been estimated that approximately 4 × 10^5^ cells enter the mature naive (FM) B cell pool daily (Srivastava et al., 2005), which is approximately 1% of the total pool size. This low rate implies that if FM B cells are maintained at roughly constant numbers, the average, net rate of turnover (the balance of loss and any self-renewal) must also be low, and indeed it has been observed that only around 50% of FM B cells are replaced over a period of 12 weeks in adults (Förster and Rajewsky, 1990, Fulcher and Basten, 1997). BrdU labelling studies have also indicated that the average lifespan of mature B cells increases with age, an effect which acts to compensate for the decrease in production (Kline et al., 1999, Fulcher and Basten, 1997).

In regard to germinal centre reactions, much attention has focused on the dynamics associated with affinity maturation, the factors influencing transitions between light and dark zones and the associated processes of proliferation and differentiation (see, for example, Victora and Mesin (2014) and Mesin et al. (2016)). However, less is known regarding the population dynamics and rules of replacement of GC B cells over extended timescales; in particular it is unclear how division and death combine to define the longevity of GC reactions, and whether their population dynamics are sensitive to host age.

Quantifying these dynamic balancing acts is important not only for understanding how to treat dysregulation of B cell homeostasis, but also for understanding how B cell repertoires evolve over time. In particular, one must be careful to distinguish between the lifetime of cells themselves, and the persistence of populations that undergo self-renewal. Collectively the progeny of naive a B cell (which comprise a B cell lineage, that we loosely refer to as a ‘clone’) may potentially persist much longer than any particular individual cell. This clonal lifetime is the pertinent quantity when measuring the persistence of antigen-specific B cell populations. Blocking B cell development by interfering with IL-7 signalling or with inducible deletion of Rag2 has indicated that mature B cell populations can persist without influx for weeks to months (Grabstein et al., 1993, Hao and Rajewsky, 2001), which defines the pool-averaged clonal lifetime.

Insights from BrdU labelling studies maybe limited due to its toxicity in the longer term and potential spatial heterogeneity in the efficiency of uptake. Also, the use of irradiated chimeras to monitor the dynamics of repopulation and maintenance is complicated by the lymphopenic environment that induces transitional cells to undergo homeostatic proliferation (Meyer-Bahlburg et al., 2008). Further, making more precise inferences regarding cell population dynamics from BrdU labelling experiments requires the use of mathematical models, and estimates of key quantities such as division and turnover (loss) rates can be sensitive to the assumptions encoded in these models (De Boer et al., 2003, De Boer and Perelson, 2013). For instance, labelling curves are often multi-phasic, indicative of heterogeneity in rates of proliferation, but fully resolving and quantifying this heterogeneity can be difficult. It can also be problematic to distinguish labelling derived from proliferation within a cell subset and from the influx of labelled cells from a precursor population, and to distinguish between potential precursor populations. To address all of these issues, we employed the method of temporal fate mapping (Hogan et al., 2015) to characterise the population dynamics and the rates and extents of tonic reconstitution of FM and naturally-occurring GC B cell compartments in healthy mice. We studied the kinetics by which new B cells percolate into peripheral subsets, and paired this information with measures of proliferation (Ki67 expression), accounting for the possible persistence in its expression across stages of development. We then confronted these data with an array of candidate mathematical models, to identify the most concise and robust descriptions of the ontogeny and dynamics of FM and GC B cells over almost the full extent of the mouse lifespan.

## Results

### Busulfan treatment permits reconstitution of the bone marrow HSC niche without perturbing peripheral mature B cell compartments

To study the dynamics of FM and GC B cells, we used a previously published method of tracking lymphocyte development in healthy mice (Hogan et al., 2015). Briefly, treatment with optimised doses of the transplant conditioning drug busulfan ablates the host haematopoetic stem cell (HSC) compartment but has no impact on mature peripheral haematopoetic lineages. We then transfer congenically labelled HSC progenitors from donor BM, which rapidly reconstitute the depleted host HSC niche. This procedure typically achieves 60-95% replacement of HSC. We then follow the replacement of mature peripheral haematopoetic compartments by the progeny of donor HSC for up to 18 months post-BMT. The kinetics of the infiltration of donor lymphocytes are rich in information regarding differentiation pathways, the fluxes between subsets and the net rates of loss within each, and the rules of replacement (Hogan et al., 2015, Gossel et al., 2017, Hogan et al., 2019).

Specifically, we generated chimeras by conditioning CD45.1 C57Bl6/J hosts with busulfan and reconstituting HSC with T- and B-cell depleted bone marrow (BM) from CD45.2 C57Bl6/J donors (Materials & Methods). To help us evaluate any influence of host age on B cell maintenance, we generated chimeras using hosts of varying ages, partioned into three groups: < 8 weeks, 8-12 weeks and ages 12 weeks or older. Previously we have shown that busulfan treatment has no detectable impact upon the long-term survival, proliferation or maintenance of peripheral T cell compartments or their progenitors (Gossel et al., 2017, Hogan et al., 2015). To confirm this was also true for B cells, we compared peripheral B cell subsets (Fig. 1A) in busulfan-treated mice with age-matched WT controls at different times following BM reconstitution. We saw no significant differences in total numbers of transitional, FM or GC B cells (Fig. 1B) or in their levels of expression of Ki67, a marker of recent division (Fig. 1C), in either spleen or lymph nodes in the weeks and months following BMT.

**Fig. 1.**
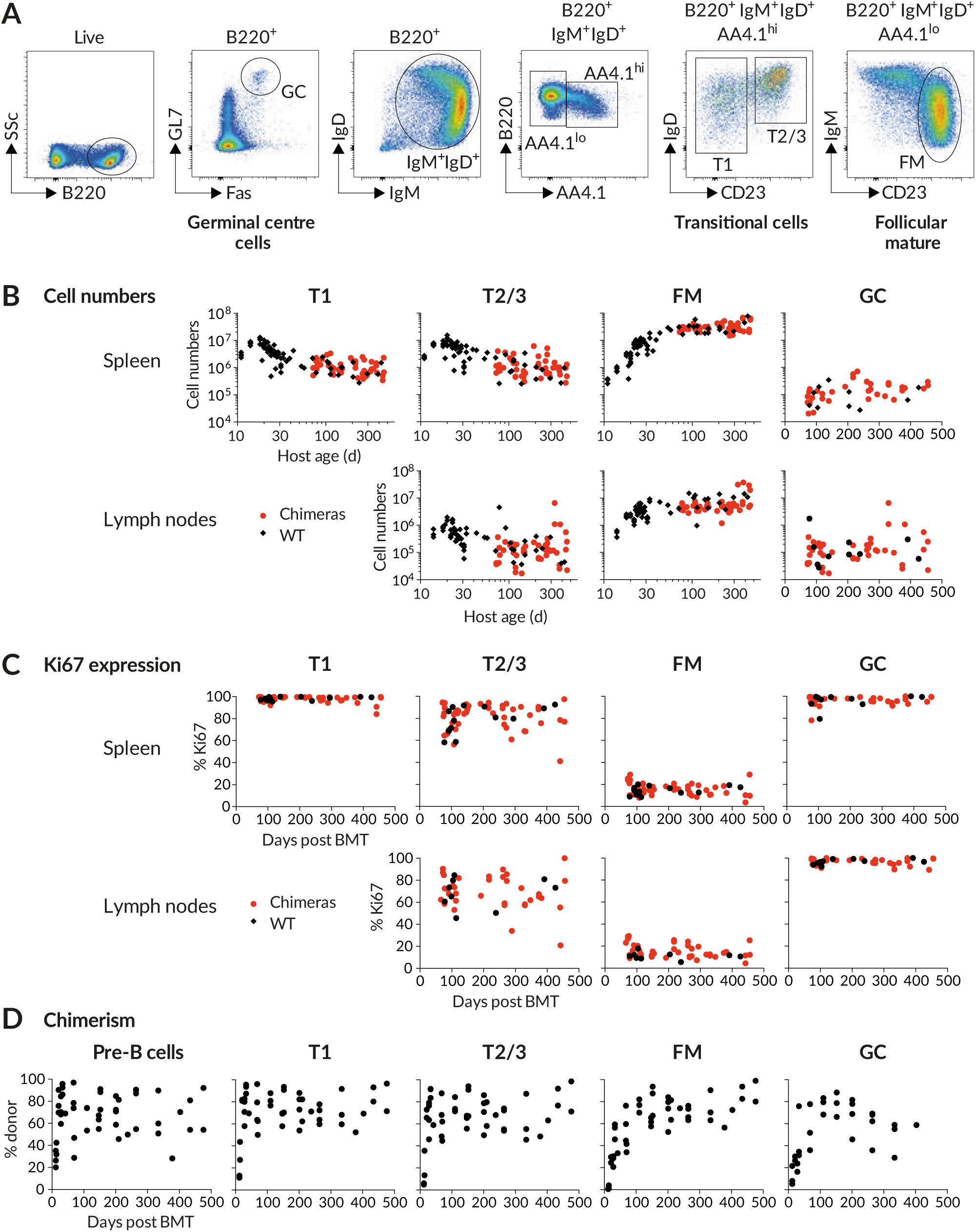
Busulfan chimeric mice exhibit normal peripheral B cell compartments. **(A)** Gating strategy to identify transitional, follicular mature and germinal centre B cells. **(B)** Comparing the sizes of B cell subsets in WT control mice and busulfan chimeras. **(C)** Comparing levels of proliferation in WT and busulfan chimeric mice, using Ki67 expression. **(D)** Host-derived B cells are gradually replaced by donor-derived cells over time. Scatter derives largely from variation in levels of stable bone marrow chimerism achieved in treated mice.

To account for variation between busulfan-treated mice in the extent of HSC replacement after BMT, we used the mouse-specific values of the chimerism among equilibrated progenitor populations to normalise the levels of donor cell infiltration into downstream peripheral B cell subsets. With this approach, a normalised chimerism of 1 indicates that a subset has attained the chimerism of its ancestral population, meaning that it has turned over completely. Previous studies of T cell homeostasis in busulfan chimeras normalised peripheral donor chimerism against equilibrated chimerism in thymic progenitors (Hogan et al., 2015). However, a similar approach normalising against donor chimerism amongst bone marrow B cell progenitors was not possible, because we observed substantial variation in donor engraftment between different bones in the same mouse (Fig. S1A). No single site was therefore representative of the entire BM compartment. However, all developing B cells migrate from BM to the spleen to continue development as AA4.1^+^IgM^hi^CD23^lo^ (transitional) cells. Since this obligate stage integrates input from all BM sites, we used the chimerism amongst these cells as a proxy for the chimerism across total BM progenitors. Following BMT, we observed a smooth transition from exclusively host-derived to donor-enriched T1 cells over the first few weeks following drug treatment, which occurred considerably more rapidly the emergence of chimerism among FM or GC B cells (Fig. 1D) but was similar in kinetics and magnitude to the emergence of chimerism among BM pre-B cells. Importantly, we then saw no trend in donor chimerism a among pre-B or T1 cells across animals over time (Fig. 1D), suggesting that BM chimerism, once established, was stable, consistent with our previous studies (Hogan et al., 2015).

### Temporal fate mapping reveals extensive replacement of mature B cell compartments by HSC progeny following BMT

We observed a high degree of correlation in donor chimerism amongst CD23^hi^ transitional 2/3 (T2/3) and FM B cells in LN, spleen and bone marrow in each mouse (Fig. 2A), confirming that these mature B cell populations are freely recirculating between the different lymphoid sites. As anticipated, the T2/3 compartment underwent complete and rapid replacement within a few weeks following BMT (Fig. 2B).

**Fig. 2.**
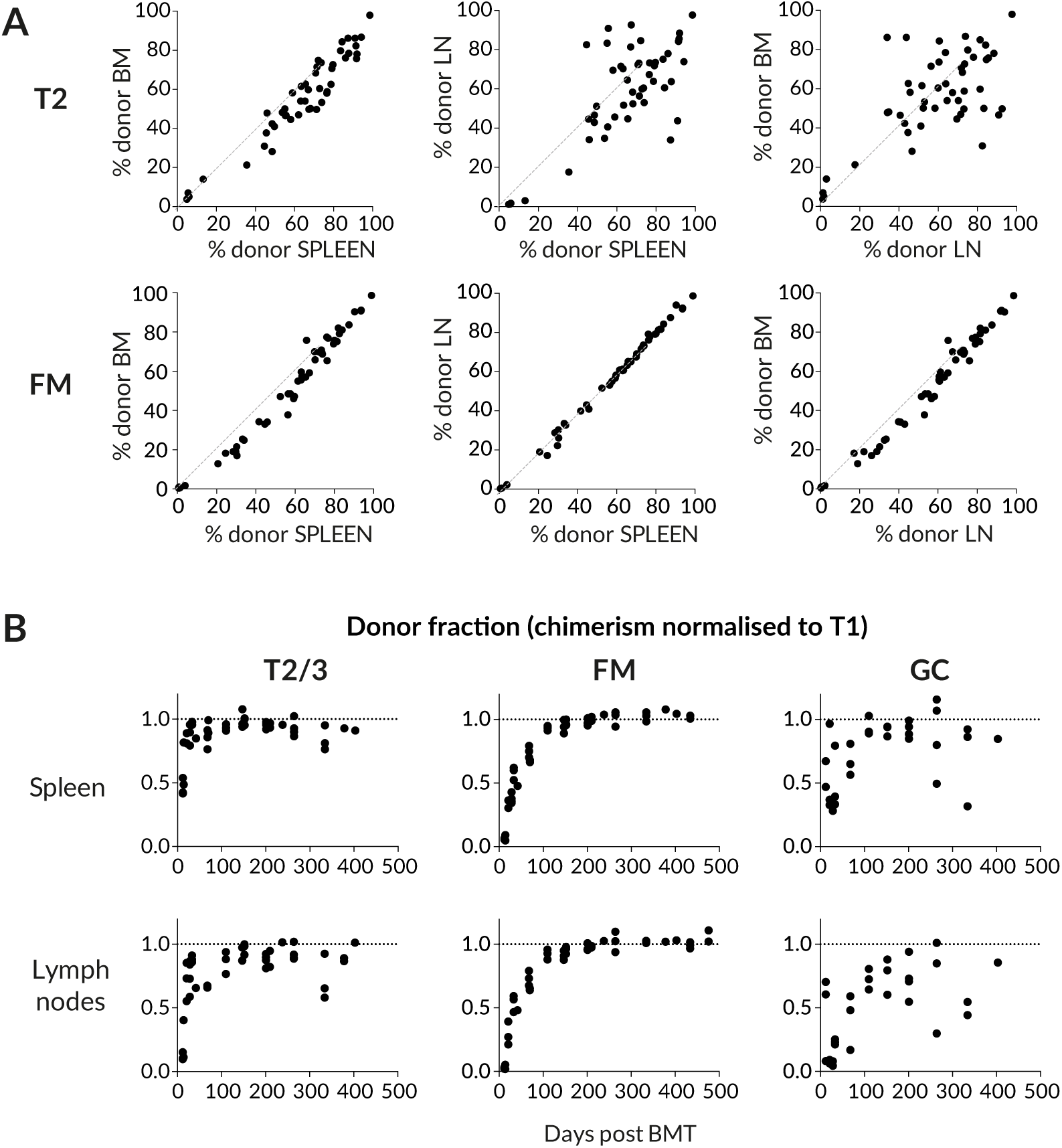
Spatial distribution and dynamics of chimerism in B cell subsets. **(A)** Comparing chimerism in the T2 and FM B subsets across bone marrow (BM), spleen and pooled lymph nodes (LN). **(B)** The kinetics of accumulation of donor-derived cells in B cell susbets. The donor fraction in each subset is normalised to that in the upstream T1 subset, to remove the effect of variation in the level of stable BM chimerism achieved across animals.

The replacement kinetics of GC B cells in spleen and lymph nodes were somewhat more noisy (Fig. 2B), but there appeared to be a more rapid replacement of within splenic GC than in lymph nodes. Since GC B cells are non-recirculating, we assumed that these two locations contained independent populations of activated B cells. Indeed, analysing individual LN separately revealed considerable variation in chimerism among GC B cells within a single host (Fig. S1B), suggestive of a degree of stochasticity with which donor or host cells are recruited to GC reactions. Nevertheless, for subsequent analyses we pooled LNs in order to measure overall levels of donor infiltration, but treated LN and spleen separately. We reasoned that the origin of stimuli driving GC formation in LN and spleen, deriving from tissue drainage and blood respectively, might result in qualitatively distinct responses.

### Quantifying cell production: Ki67 expression reflects self-renewal but is also inherited across stages of B cell development

To examine the role of proliferation in B cell development, we also measured levels of Ki67, a nuclear protein that serves as a marker of recent cell division. In T cells, Ki67 expression is induced at the G1 stage of the cell cycle and persists for more than three days after mitosis (Gossel et al., 2017, Hogan et al., 2013). Similar decay kinetics have been described in human mammary epithelial cell lines *in vitro* (Miller et al., 2018).

Analysis of Ki67 expression revealed variation across stages of B cell development in bone marrow, transitional stages and mature B cells (Fig. 3A). Pre-B cells and GC B cells both undergo extensive proliferation and exhibit a high, unimodal distribution of Ki67 expression. It was also readily detectable amongst transitional B cells, but was expressed by only a small subset of FM B cells (Fig. 3A). It has been shown that transitional populations in the spleen do not divide as they mature, and that following BrdU administration, labelled T2 cells are detected in the spleen within 2 days (Srivastava et al., 2005), a maturation time that is shorter than the lifetime of Ki67. Therefore, Ki67 seen in transitional populations is likely residual expression from division events among precursors in the bone marrow. Consistent with this inference, Ki67 abundance in transitional populations remains broadly unimodal in distribution but its median level falls as they mature into FM B cells (Fig. 3A). Given the short transit time through the transitional stages, it is possible that the low level of Ki67 expression in FM B cells (Fig. 1B and Fig. 3A) could also derive at least in part from recently divided BM precursors, as well as from self-renewal. Evidence for inheritance of Ki67 in the FM pool is that we saw a transient elevation in the frequency of Ki67^hi^ cells in donor cells over that in host cells soon after BMT, but levels declined and converged with host cell levels after ∼100 days (Fig. 3B). We infer that this behaviour derives from the difference in the mean ages of donor and host B cells, which is more pronounced soon after BMT when all donor-derived FM cells have recently entered the compartment, and not from any intrinsic differences in host and donor cell behaviour. Therefore, in the modelling analyses described below we assumed that Ki67 expression within each B cell subset could derive from division and/or the influx of Ki67^hi^ progenitors.

**Fig. 3.**
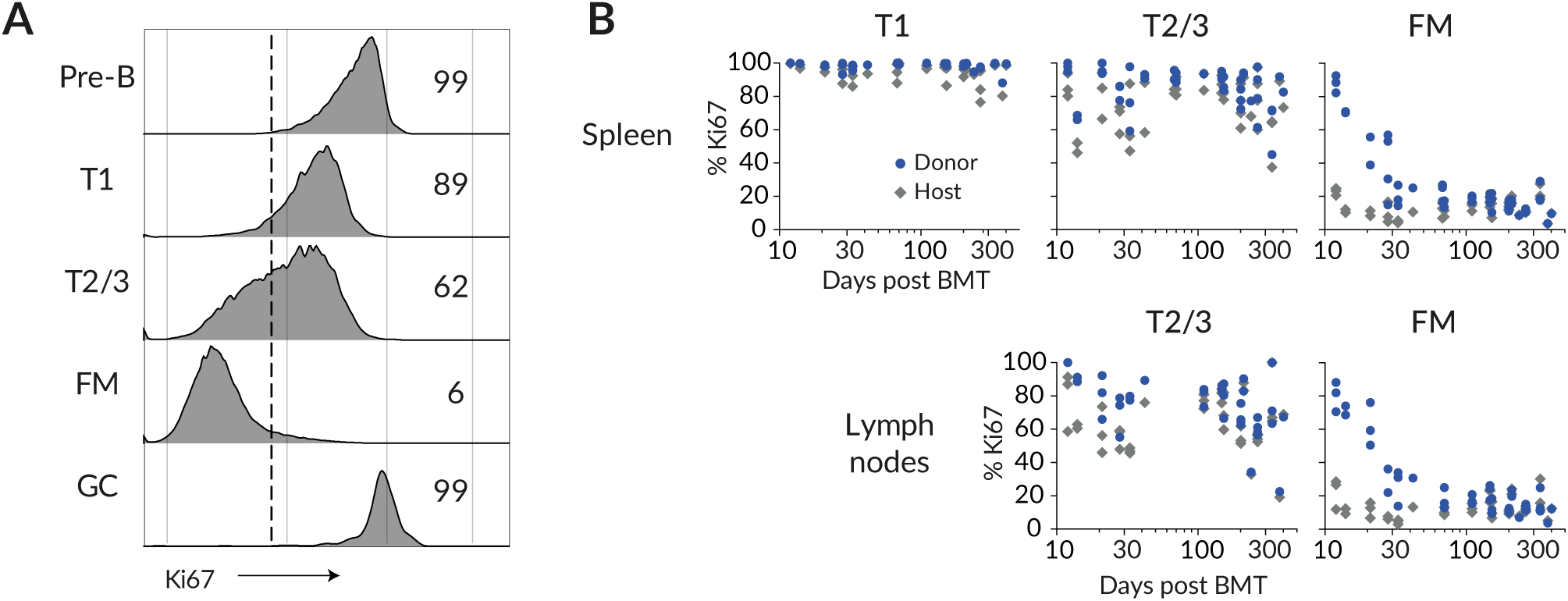
Levels of proliferation vary throughout B cell development. **(A)** Ki67 expression in different B cell subsets; Pre-B cells in bone marrow, transitional T1/2/3 cells in spleen, and FM and GC B cells in spleen. **(B)** Ki67 frequencies amongst donor- and host-derived B cell subsets with time post BMT.

### Follicular Mature B cells are a homogeneous, slowly dividing population whose residence time increases with age

Given the time-varying fluxes of donor cells through multiple stages of development, extracting the maximum information from these timecourses requires mathematical modelling. We have previously used this approach using busulfan chimeric mice to quantify the developmental and homeostatic dynamics of naive CD4 and CD8 T cells (Hogan et al., 2015) and memory CD4 T cells (Gossel et al., 2017, Hogan et al., 2019).

We began by studying FM B cells. We assumed that they recirculate freely between lymph nodes and spleen, given the close similarity in chimerism in the two compartments (Fig. 2A) and indeed across all lymphoid organs (Fig. S1B). Therefore we pooled the numbers of FM B cells recovered from spleen and lymph nodes and assumed they follow the same dynamics in each. We attempted to describe these dynamics in mice aged between 70 and 600 days with a variety of mathematical models (Fig. 4A). In each, we assumed newly-differentiated FM B cells are generated at a rate proportional to the size of their precursor population, which was assumed to be either T1, T2, or T1 and T2 combined. Describing the timecourses of these ‘source’ populations with empirical functions (Fig. S2 and Text S4), we then aimed to identify the combination of model and precursor population that best described FM B cell dynamics. The simplest model (Fig. 4A, top) assumed that FM B cells, whether host or donor, are generated from their precursors at the same constant *per capita* rate, and form a homogeneous population which undergoes turnover (loss) and self-renews through division, both at constant *per capita* rates. This model predicts a smooth, continuous approach to stable chimerism of FM B cells with eventual complete (and repeated) replacement. However, the precise shape of this curve is rich in information regarding the processes of influx and loss. To test for any more complex homeostatic dynamics, we considered four extensions to this basic model. In the first, the rates of turnover or division might vary with host age (the ‘time-dependent turnover’ or ‘time-dependent division’ models). In the second, the FM B cells are assumed to be homogeneous with constant rates of turnover and division, but are fed from transitional B cells at a *per capita* rate that changes with age (‘time-dependent recruitment’). In the third extension, FM B cells comprise two independent subpopulations turning over at different rates (‘kinetic heterogeneity’). In this scenario the donor chimerism will initially increase rapidly as the subpopulation with faster turnover is replaced, followed by a more gradual approach to stable chimerism as the more persistent subpopulation, with slower turnover, is replaced. In the fourth extension (the ‘incumbent’ model), we allowed for the possibility that a population of host-derived cells established early in life remains stable in numbers and is not replaced by cells recruited later in life. Such a model allows for less-than-complete turnover, or a normalised chimerism stabilising at a value less than 1. See Text S1 for details of the mathematical formulation of the models.

**Fig. 4.**
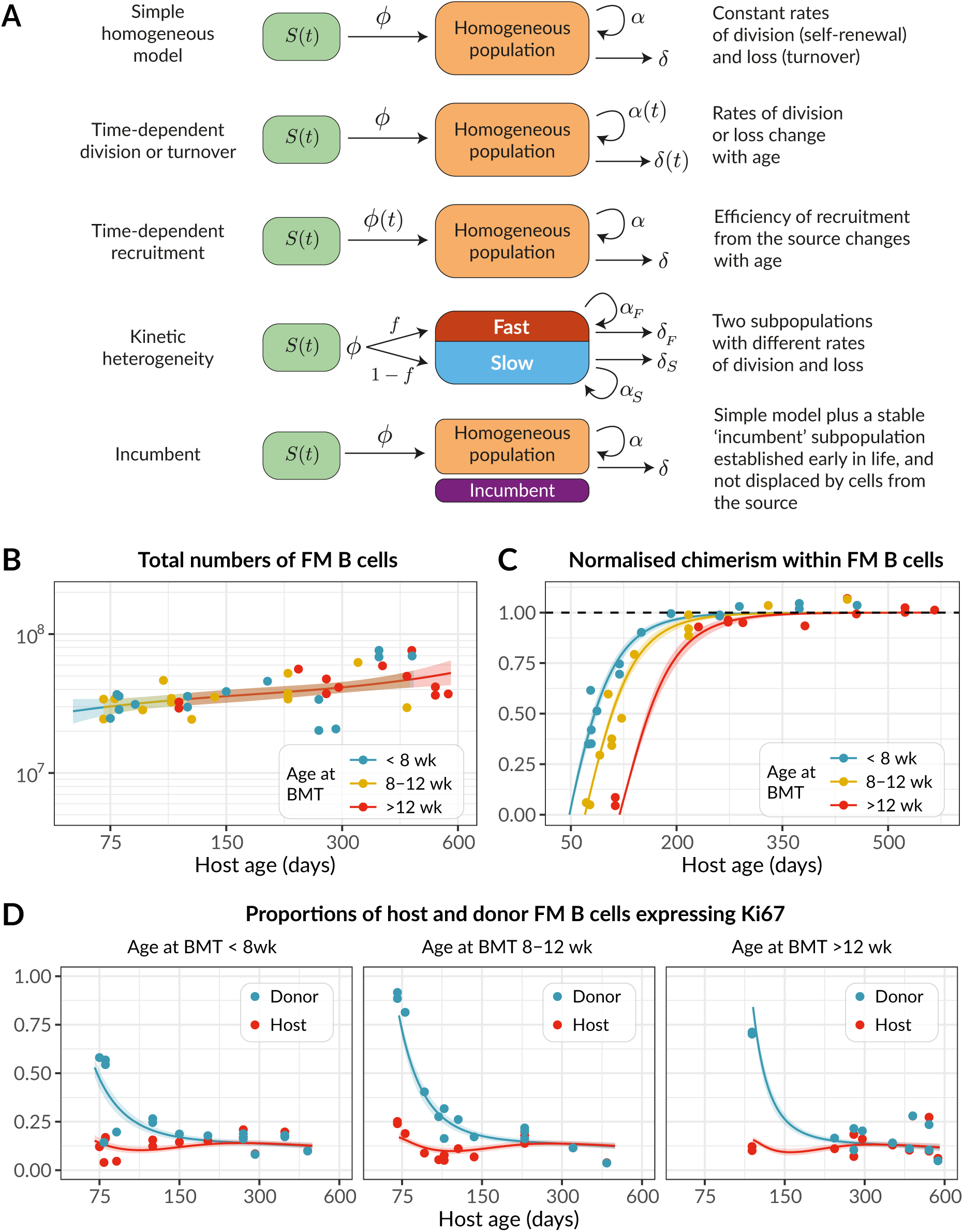
Modelling the population dynamics of FM B cells in busulfan chimeric mice. **(A)** Schematics of the candidate mathematical models of FM B cells (orange) fed by a precursor population (green). **Panels B-D:** Observed (points) and best-fit values (lines) of **(B)** total FM B cell numbers, from pooled spleen and lymph nodes; **(C)** their chimerism, normalised to that in T1; and **(D)** the proportions of host and donor FM B cells expressing Ki67. Lines show the predictions from the best-fitting (time-dependent turnover) model, in which the mean residence time of FM B cells increases with mouse age. Colours in panels B and C denote groups of mice who underwent BMT at different ages. Shaded regions are prediction intervals, generated by drawing samples from the posterior distribution of parameter estimates and plotting the 4.5 and 95.5 percentiles of the resulting model predictions. Fitted values were specific to each mouse, with its particular age at BMT; model predictions shown here were generated using the mean age at BMT within each group.

The kinetic of replacement of existing cells with immigrant cells is determined primarily by the average net rate of loss – the balance of cell death, any onward differentiation, and proliferative renewal (Text S2). We refer to the inverse of the net loss rate as the clonal lifespan; it measures the persistence of a population of B cells that is subject to both loss and a degree of self-renewal. The clonal lifespan may be much longer than the expected time any one cell spends within that population before it dies or differentiates, which we refer to as the residence time.

To estimate parameters and compare the support for the models, each was fitted simultaneously to the timecourses of FM B cell numbers, the chimerism within FM B cells normalised to that in T1 (the earliest common precursor to all populations considered), and the proportions of host and donor FM B cells expressing Ki67 (Fig. 4A-C). Including the Ki67 expression in donor and host populations allowed us to resolve the net loss rate into rates of death/differentiation (yielding the residence time) and proliferation (yielding the interdivision time). Our approach to fitting is outlined in Materials & Methods and is described in detail in Text S3.

We could immediately reject the incumbent model because the chimerism of the FM B cell compartment reached that of T1 cells (Fig. 4B), indicating no evidence for a persistent host-derived FM B cell population. We found the strongest relative support (64%, Table S1) for the model in which the rate of turnover (loss) of FM B cells declined with host age but their rate of division remained constant, and with T1 cells as their direct precursor. Fits are shown as solid lines in Fig. 4A-C, and parameter estimates are in Table 1.

**Table 1.**
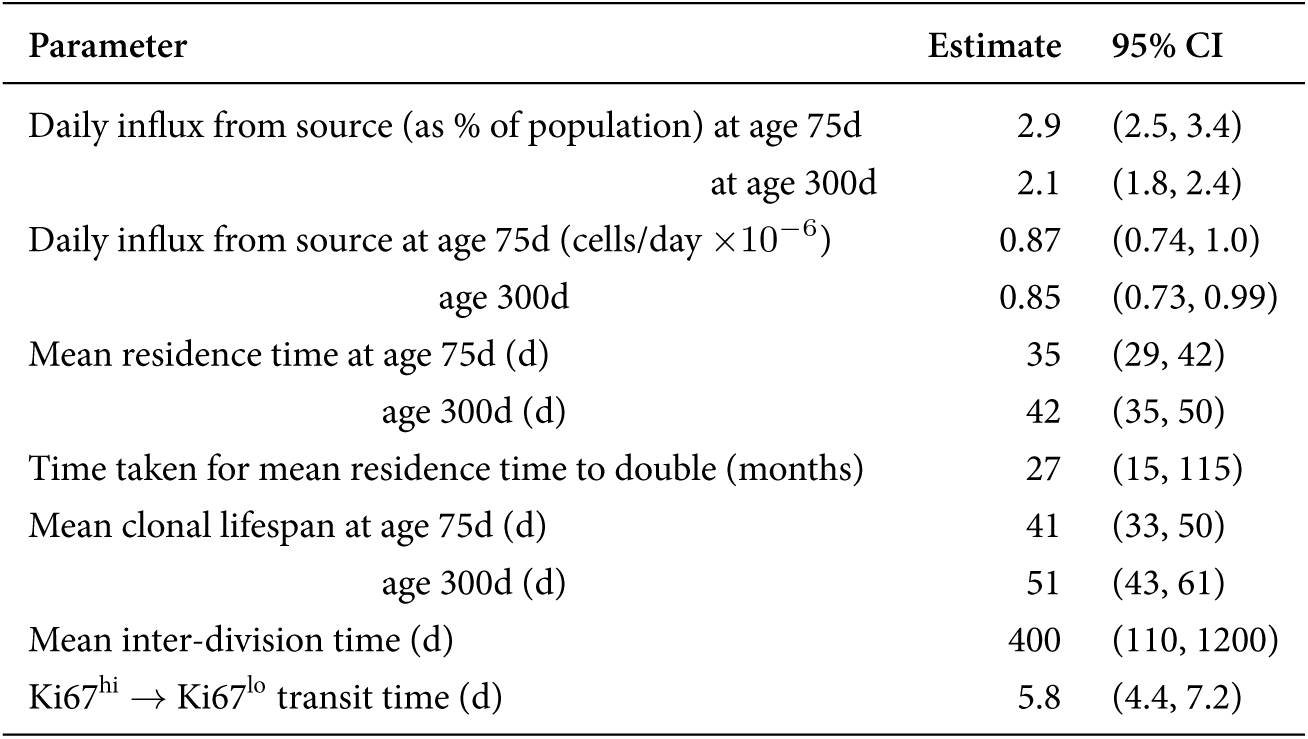
Parameter estimates from the best-fitting model of FM B cell homeostasis. 95% credible intervals were estimated by taking the 2.5 and 97.5 percentiles of the posterior probability distributions of the parameter values.

We estimate a steady daily influx of around 0.9 × 10^6^ new FM B cells per day from T1 precursors into the spleen and LN of mice aged between 75d and 500d, deriving from the assumption of a constant *per capita* rate of recruitment and the relatively stable numbers of T1 precursors in this age range (Fig. 1A). This flux is almost double an estimate of the number of new FM B cells entering the spleen daily in 56d-old mice (Srivastava et al., 2005). We infer that FM B cells have a mean residence time of roughly 5 weeks in 75 day-old mice, and that this increases slowly over time (6 weeks at age 300 days and almost 9 weeks at age 2 years). These estimates are in line with those from older studies of BrdU labelling among mature B220^hi^ HSA^low^ B cells, which are predominantly FM cells (Förster and Rajewsky, 1990, Fulcher and Basten, 1997).

While in adult mice approximately 10% of FM B cells express Ki67 (Figs. 1B and 4D), we infer that this level of expression derives almost entirely from newly generated FM cells who inherit it from their pre-transitional, highly proliferative bone marrow precursors. As described above, this conclusion derives largely from the observation that donor-derived FM B cells, which soon after BMT are highly enriched for newly generated cells, transiently exhibit significantly higher levels of Ki67 than the more established host cells (Fig. 4D). We infer that FM B cells themselves divide rarely – roughly once a year, though this estimate comes with some uncertainty. Because this self-renewal is slow, the average clonal lifetime is only slightly longer than the mean residence time of individual cells themselves. Therefore, the naive FM B cell compartment in adult mice relies almost entirely on the influx of new cells – and is therefore constantly supplied with new receptor specificities – for its maintenance throughout life.

### Developmental dynamics of FM B cells differ in young and adult mice

Next, we studied the accumulation of FM B cells early in life to understand how the dynamics of their establishment in lymphoid organs compares to their dynamics in adult mice. Their T1 precursors dramatically increased in number up to age 20 days, declined continuously for a further 20-30 days, and were maintained stably thereafter (Fig. 1A). Correspondingly, FM B cell numbers increase rapidly up to age 30 days, followed by the much slower but persistent increase that we modelled in adults (Fig. 1A). We wanted to explore whether the processes of generation and maintenance of FM B cells from T1 precursors that we characterised in adults followed the same dynamics early in life. To do this, we used our best-fit time-dependent turnover model and its parameter estimates from adult mice together with the timecourse of T1 B cell numbers in young mice (Fig. 5A; see Text S5 for details of its parameterisation) to predict the kinetics of accumulation of FM B cells from age 10 days onwards, extrapolating the exponentially-decaying loss rate back to the earliest timepoint (age 14 days). We found that the predicted FM B cell numbers substantially overshot the observations (∼3-fold higher at age 4 weeks; Fig. 5B).

**Fig. 5.**
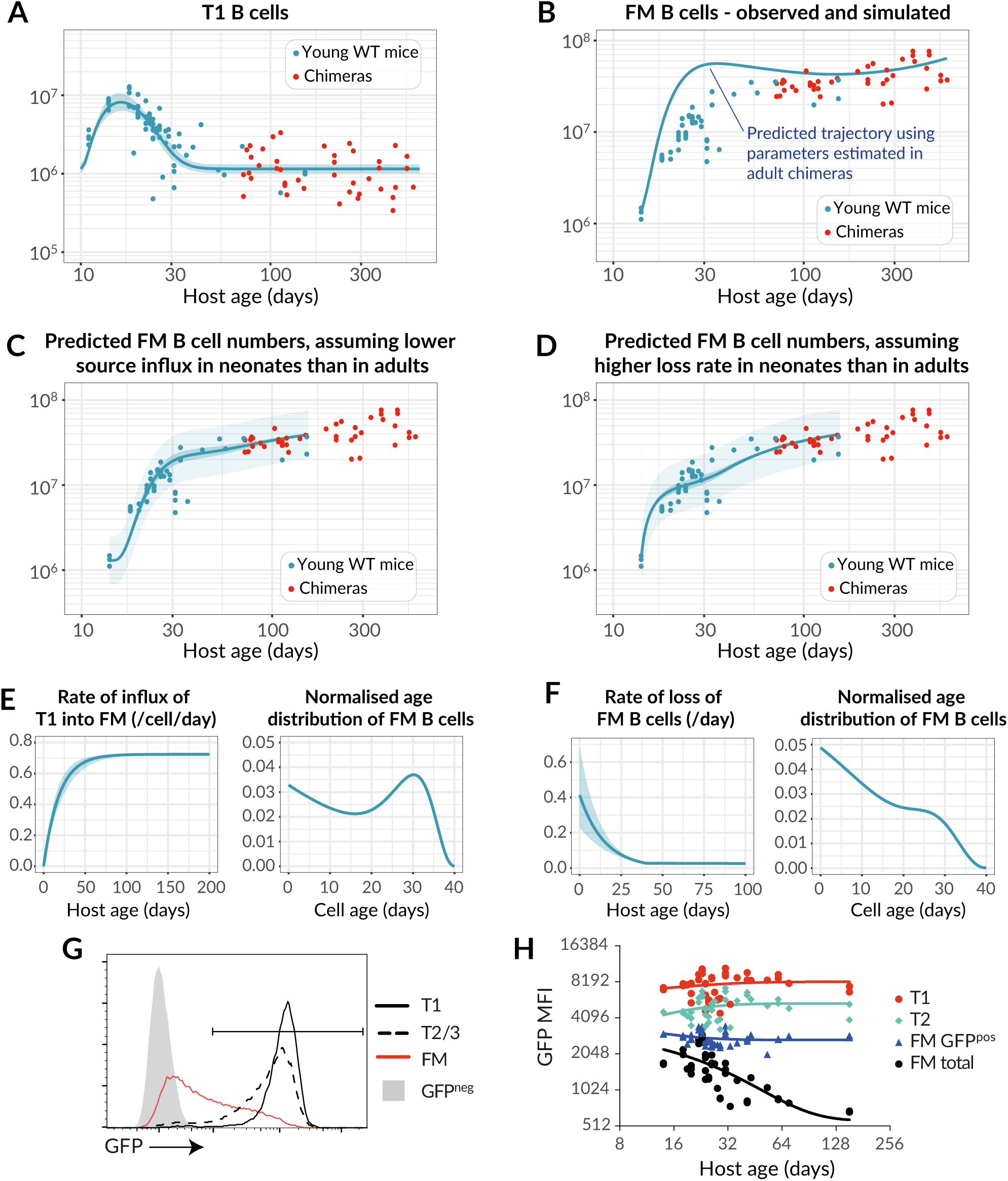
Establishment of the FM B cell compartment in young mice is characterised by progressively increasing recruitment from T1 precursors with age. **(A)** Observed numbers of splenic T1 B cells recovered from both young WT control mice (blue) and adult busulfan chimeras (red), described with a single fitted empirical function (see Text S5). **(B)** Total numbers of FM B cells recovered from the same mice. The blue line shows the accrual of FM B cells from age 14d predicted by the best-fitting model of of FM B cell dynamics in adult chimeras (Fig. 4), using the timecourse of T1 precursors in young WT mice. **(C-D)** Fits to the numbers of FM B cells in young WT mice (blue dots), using extensions of the best-fitting model in which either the rate of recruitment from the T1 pool increases with age in young mice (C), or the death rate of FM cells decreases with age (D). **(E)** Best fitting functional form for the change in *per capita* daily rate of influx from T1 to FM, and the predicted age distribution of FM B cells in 40d-old mice. **(F)** Best fitting functional form for the change in loss rate of FM with age, and their predicted age distribution in 40d-old mice. **(G)** GFP levels of subsets of splenic B cells in 6wk-old Rag2-GFP mice. **(H)** Variation in mean fluorescence of GFP in B cell subsets in Rag2-GFP mice, with age.

This mismatch indicated that either (i) cells flow from T1 to the FM B cell pool at a lower *per capita* rate early in life; and/or (ii) FM B cells in young mice are lost much more rapidly than those in adult mice, at even greater levels than predicted by the best-fitting model of age-dependent loss. We tested these two hypotheses in silico by allowing the *per capita* rate of influx from T1 to increase progressively with host age, or by augmenting the death rate of FM B cells early in life (see Text S5). We then fitted these two models to the FM B cell counts in young mice (Fig. 5C and D). Both of these extensions described the data well, but they made very distinct predictions regarding the age distribution of FM B cells in young mice (see Text S5 for details). Increasing recruitment from T1 predicted a broad distribution of cell ages (Fig. 5E), while higher loss rates in young mice predicted a preponderance of younger cells (Fig. 5F). To test these predictions, we analysed FM B cell development in Rag2-GFP transgenic mice. In these mice, GFP expression is induced in bone marrow progenitors during RAG mediated BCR recombination, and persists into peripheral transitional and mature FM B cell populations (Fig. 5G). We could then use the distribution of GFP expression within a population as a surrogate of its age distribution. As expected, GFP levels in the T1 and T2/3 compartments, which turn over rapidly, were uniformly high and did not vary with host age (Fig. 5H). Average GFP fluorescence in FM B cells was high in young mice, and as expected declined with age, as mature GFP-negative cells accumulated. Significantly, however, the average GFP expression in GFP-positive FM B cells, which are newly generated, was also invariant with host age. If newly generated FM B cells were shorter-lived in neonates, we would expect a relative enrichment of GFP^hi^ FM B cells in younger mice, with an associated higher population-average GFP expression than in adults. This was not observed. Therefore, the relatively low rate of accumulation of FM B cells in neonates most likely derives largely from a lower rate of differentiation from T1 progenitor cells early in life, rather than shorter lifespans of FM B cells.

### Germinal centre B cells in spleen and lymph nodes exhibit distinct dynamics

We next applied a similar modelling approach to examine the dynamics of naturally occurring GC reactions in naive mice throughout life. Although the stimuli that drive formation of these GC reactions have not been characterised, analysis of germ free mice revealed similar numbers of GC B cells in spleen but reduced numbers in LNs as compared with WT controls in conventional facilities (Fig. S3). These observations indicate that naturally occurring GC reactions are driven by a self/endogenous stimuli in the spleen and a more dominant foreign source of antigen in LNs.

The number of GC B cells in the spleen gradually increased with age (Fig. 6A), implying either a gradual increase in the rate of influx from their precursors, and/or increases in GC B cell lifespan or proliferation rate with age. The chimerism of splenic GC B cells stabilised within ∼100 days (Fig. 6B), more rapidly than that of FM B cells (∼150 days, Fig. 4C). This asynchrony in development discounts FM B cells as the precursors of splenic GC B cells. Therefore, we inferred that splenic GC B cells derive directly from immature transitional B cell subsets. We then fitted the models illustrated in Fig. 4A to the timecourses of numbers, chimerism, and Ki67 expression of splenic GC B cells. However, all of the models received comparable levels of statistical support (Table S2, rows shaded in grey) preventing us from clearly discriminating between them.

**Fig. 6.**
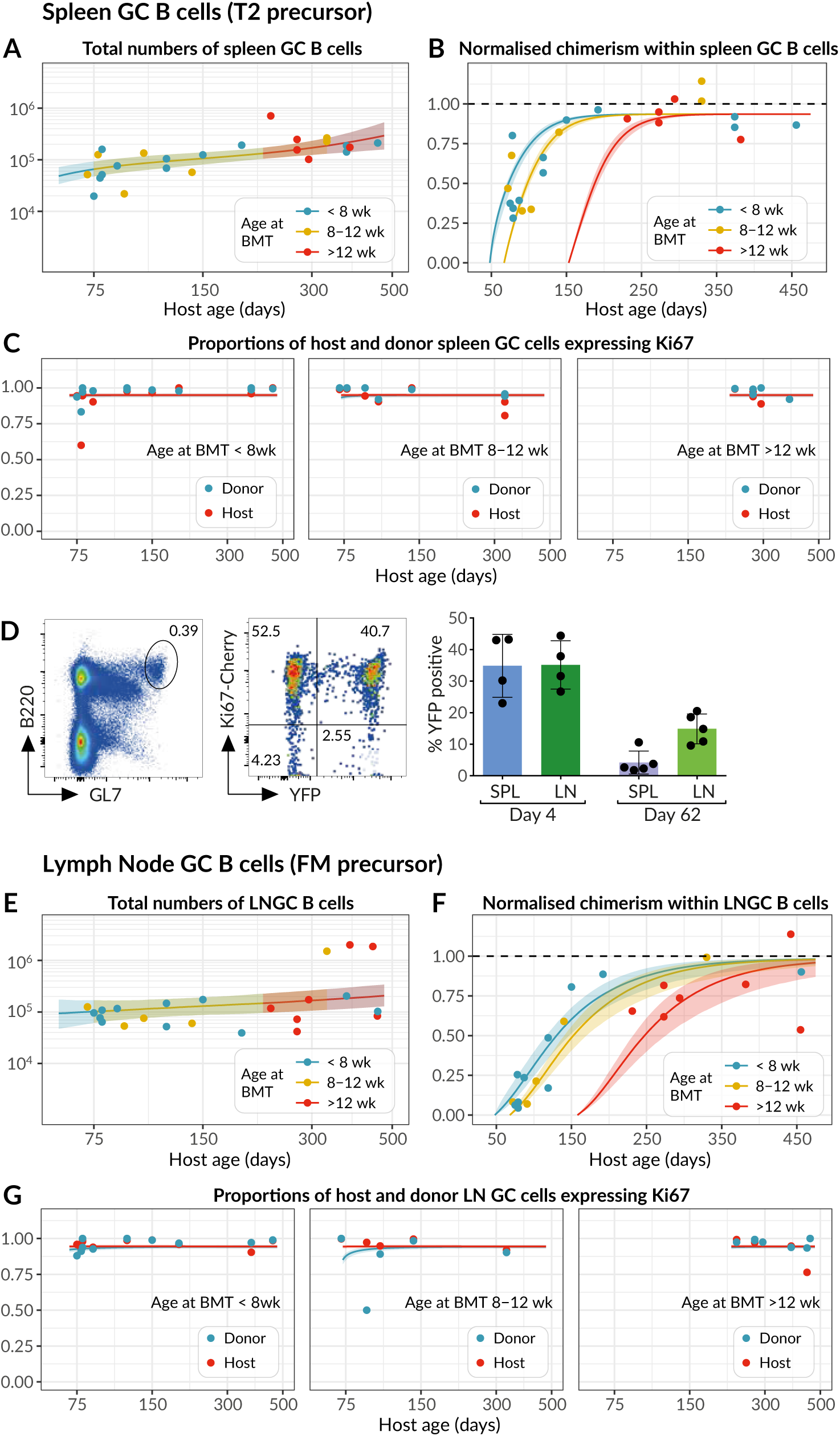
Modelling the population dynamics of germinal centre B cells in spleen and lymph nodes. **(A-C)** Splenic GC B cell dynamics, with solid lines denoting the best fits to total cell counts, normalised chimerism and Ki67 expression stratified by host and donor cells. Shaded regions are prediction intervals. described in Fig. 4. Line colours indicate grouping by age at BMT. Curves were generated using the mean age at BMT within each group. **(D)** Using the Ki67-YFP reporter mouse to track cohorts of divided cells entering the GC pools in spleen and lymph nodes. (Left panel) Gating strategy for GC B cells. (Middle panel) Cells divided during tamoxifen administration express Ki67-Cherry transiently and YFP heritably. (Right panel) Frequencies of YFP^+^ cells among GC B cells decrease with time as they are diluted by YFP^−^ immigrant cells. **(E-G)** Analogous fits to the dynamics of lymph-node GC (LNGC) B cells.

This uncertainty stems from the relatively noisy approach to stable chimerism among GC B cells, which rather poorly constrains their net loss rate; and the nearly saturating and constant levels of expression of Ki67 among host and donor cells (Fig. 6C), which then provide relatively little information regarding the levels of inheritance of Ki67 from the source and the division rate of GC B cells themselves. To increase our ability to discriminate between models we exploited this high level of Ki67 expression. We generated a novel fate mapping mouse strain in which an inducible CreERT2 construct was expressed from the endogenous *Mki67* locus, alongside a Ki67-Cherry fusion protein. Crossing these with *Rosa26*^*RYFP*^ Cre reporter mice generated a strain in which, following induction of Cre activity by the inducer tamoxifen, dividing cells and their progeny could be indelibly labelled by expression of YFP. Treating these Ki67 reporter mice with tamoxifen for just 4 days resulted in labelling of a substantial and similar fraction of GC B cells in both spleen and lymph nodes, which declined 8 weeks after induction (Fig. 6D). The fold reduction in YFP expression allowed us to place a tighter prior on the net loss rate of GC B cells (Text S6). Re-fitting the models using this information then revealed the strongest support for the model of splenic GC as a kinetically homogeneous population, fed by T2 B cells at a rate that increases gradually with mouse age (57% of the model weights, Table S2; fits shown in Fig. 6A-C).

As expected, we inferred that splenic GC B cells are more dynamic than FM B cells, with mean cell lifetimes and mean inter-division times both roughly 12h. Remarkably, the net effect of these processes yields a mean clonal lifespan of about 30 days (Table 2); thus, B cell lineages in GC are preserved for several weeks with very rapid turnover. Because the numbers of splenic GC B cells do not change rapidly with age in adult mice and so can be considered close to equilibrium, this clonal lifespan is the main determinant of the timescale of replacement of host with donor cells (Text S2). We estimate that number of cells entering the splenic GC pool per day as a proportion of the pool size is between 4-5% in adulthood, which is indeed close to the daily loss rate (the inverse of the clonal lifespan).

**Table 2.**
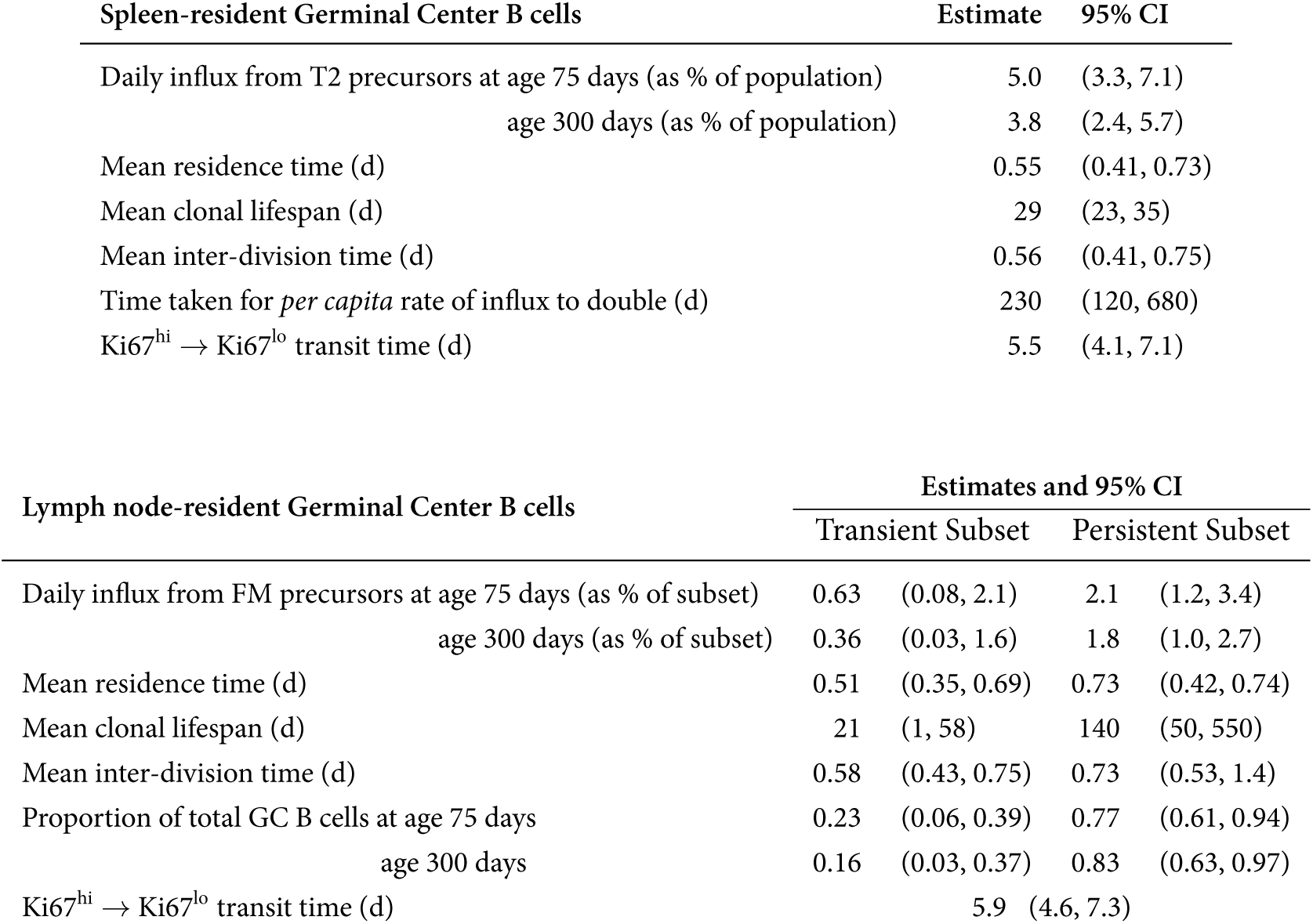
Parameters governing homeostasis of germinal centre B cells in the spleen and lymph nodes. Upper table: Estimates from the best-fitting model of splenic GC B cell dynamics, in which the rate of influx of new cells into the compartment from T2 precursors increases with host age. 95% credible intervals (CI) were derived from the 2.5 and 97.5 percentiles of the posterior distributions of the parameter values. **Lower table:** Parameter estimates for lymph node GC B cell homeostasis, derived from the best-fitting (kinetic heterogeneity) model with FM B cells as the precursor population and two populations with different rates of division and turnover. Both populations are assumed to share the same Ki67 lifetime.

We performed a similar analysis of lymph-node GC (LNGC) B cell dynamics, again exploiting the Ki67 reporter mice to place bounds on the average net loss rate of these cells and to increase our ability to distinguish between models. We found strong evidence for the existence of two subsets of LNGC B cells with distinct but rapid kinetics, with FM B cells as the direct precursor of both (84% relative weight, Table S3, unshaded rows; model fits shown in Fig. 6E-G), with lower support for a T2 precursor (11% relative weight, Table S3). However, the existence of this heterogeneity, and the visual similarity of the fits deriving from the two differentiation pathways (not shown), suggests that LNGC B cells may in fact be fed to different degrees by both FM and T2 cells.

We estimated a low rate of influx into the LNGC pool (at most 2% of the pool size per day), with a transient subset consisting of short-lived clones (lifetime∼20d) and a persistent subset with a clonal lifetime of ∼140d. The persistent clones comprise approximately 80% of LNGC B cells, and their lifespan is the key determinant of the slow rate at which donor cells infiltrate the compartment (Fig. 6F).

## Discussion

Our analysis of naive (follicular mature) B cell homeostasis revealed continuous replenishment from the bone marrow throughout life, with the entire compartment subject to replacement within 200 days, and the best model of the dynamics of this compartment was one in which all naive FM B cells follow the same rules of replacement irrespective of their residence history. Although it appears that all naive B cells are made and remain equal with respect to homeostatic fitness, we did find strong evidence that age dependent changes in host environment influenced both FM B cell maintenance and development. The most strong supported model, in which turnover (loss) decreases with time, indicated that B cells live almost twice as long in aged mice. We also found that development of new B cells from transitional precursors was relatively inefficient in neonates, despite an abundance of T1 and T2 precursors. It therefore appears that host environment is the biggest factor influencing B cell homeostasis, rather than cell-intrinsic changes. Age associated B cells (AABCs) are a subpopulation of IgM^hi^ CD23^lo^ AA4.1^-^ FAS^+^ B cells that appear in ageing hosts (Naradikian et al., 2016). We observed AABC in aged busulfan chimeras and found they were predominantly of donor origin (Fig. S4), indicating that they are generated by newer cells later in life. This observation suggests that their emergence is also associated with a changing host environment and not from a cell-intrinsic increase in the propensity of ageing cells to differentiate. In the latter case, one would expect a larger representation of host-derived cells within that population.

We also assessed the contribution of cell division to peripheral B cell homeostasis, through analysis of Ki67 expression. The extensive proliferation of germinal centre B cells is a well-recognised feature of their development, but the role of cell division in supporting naive B cell homeostasis is less clear. The persistence of Ki67 for a few days following mitosis complicates the interpretation of its expression among differentiating populations, particularly when differentiation occurs over similar or shorter timescales. Our analysis revealed that Ki67 levels in peripheral naive B cells and transitional populations derive almost exclusively from proliferating bone marrow precursors and not from proliferative self-renewal. Indeed, the best fitting model, that allowed for inheritance of Ki67 from precursors, indicated that FM B cells divide extremely rarely, if at all.

Germinal centres are readily detectable in laboratory mice, even in the absence of deliberate infection. Although the range of stimuli that elicit such responses has not been fully characterised, commensal organisms in the gut may provide a significant antigenic drive (Reboldi and Cyster, 2016). The constant flux of new cells into these structures that we observe in the busulfan chimeric mice is consistent with the observation that new FM B cells are continually recruited into chronic GC reactions (Schwickert et al., 2007, Shulman et al., 2013) or into the same GC after repeated immunisations (Bergqvist et al., 2013). The detailed dynamics of GC reactions over short timescales, typically those of acute infections, have been modelled extensively (see, for example, Kepler and Perelson (1993), Figge (2005), Anderson et al. (2009), Meyer-Hermann et al. (2012), Robert et al. (2017)), but here we focused on GC B cell production and turnover over timescales of months to years. We revealed that GC B cell lineages – which we loosely referred to here as clones, belying the process of affinity maturation – persist for many weeks. Strikingly, we were still able to resolve the remarkably rapid cellular dynamics underlying this persistence. Our inference that average individual GC B cell lifespans are very short is consistent with observations of the frequency of apoptotic cells, which led to the conclusion that least 3% of GC B cells die per hour (Wittenbrink et al., 2011). This figure translates to minimal death rate of 0.73/d or an upper bound on the mean lifespan of 1.4d, consistent with our estimates of 12-18 hours. Our estimates of interdivision times of roughly half a day are also comparable to other estimates derived from BrdU labelling (Anderson et al., 2009). The fine balance between these two rapid processes underpins the extended lifetimes of GC B cell lineages that we expose here.

Our analysis also made distinct predictions regarding GC reactions in spleen and lymph nodes. Splenic GC appeared to involve homogeneous dynamics, sourced primarily from T2/3 cells, although the data were sufficiently noisy to perhaps obscure any kinetic substructure. In contrast, we found evidence of heterogeneous kinetic substructures within GC reactions in lymph nodes, which are fed predominantly from mature FM B cells. These distinct dynamics, particularly with regard to the influx of new cells, suggest differences in the nature of antigen exposure in these organs. Indeed, GC B cells were readily detectable in the spleens of germ free mice while LN GC B cells were substantially reduced in number in the same mice. We speculate that splenic GC B cell clones are generated predominantly by weak responses to self-antigens and these reactions are fed at a relatively high rate by new B cells almost as soon as they emerge from development. In contrast, lymph node GC B cell clones are likely derived from stronger and rarer cognate reactions of FM B cells to foreign antigens draining from epithelial barriers. Notably, the subpopulations we identified in constitutive lymph node reactions were still both highly proliferative, and so probably do not correspond to the kinetically distinct populations of B cells found in light and dark zones within germinal centers (Mesin et al., 2016, Meyer-Hermann et al., 2012). This heterogeneneity could arise from multiple sources. GC reactions could be seeded from both FM and T2 sources, and they could also involve both newly-stimulated naive B cells and recirculating memory cells, which may exhibit different kinetics. Another possibility is that, rather than representing subpopulations within the same GC, the heterogeneity we detected among LN GC B cells derives in part from the pooling of multiple lymph nodes in our analysis, and indicates that different lymphoid organs exhibit different degrees of GC B cell turnover.

We found little statistical support for models in which transitional T2 B cells are the direct precursors of FM B cells, which is perhaps surprising as they represent a more advanced stage of development than T1 cells. Notably, we found a relatively weak correlation between the chimerism of T2 cells in spleen and lymph nodes within the same mouse (Fig. 2A), suggesting that T2 cells are a spatially heterogeneous population. This result suggests that a more refined modelling approach would account for the circulation of transitional and mature B cell populations between the spleen and secondary lymphoid organs, at the cost of having to estimate a greater number of free parameters.

Our study reveals the importance of continual influx to the maintenance of naive B cell compartments in mice that bears similarities to the mode of maintenance of naive T cells in adult mice (Hogan et al., 2015, Rane et al., 2018). In adulthood, both naive B and T cells have lifespans of several weeks and are reliant upon a daily influx of new cells that comprises a few percent of the pool size. We also find evidence for increased longevity of both populations as the mouse ages, that compensates to some degree for the waning of transitional B cell precursors in young adulthood, and the more substantial and longer-term involution of the thymus. The mechanisms that enhance lifespan, however, contrast between the two lymphocyte lineages. T cells exhibit cell-intrinsic adaptations such that older cells become fitter as they age and are preferentially retained in the repertoire (Rane et al., 2018), while we infer that increased B cell longevity is achieved by changes in the host that impact B cell populations uniformly. Consequently, B cells retain their homogenous homeostatic properties, while T cell compartments become increasingly heterogeneous with age, with evidence of naive T cell clones being retained for many months and even years (Hogan et al., 2015). The cell-intrinsic adaptation of T cells may be driven by the self-MHC recognition that has been shown to be essential for their long term survival (Martin et al., 2006). The environmental factors responsible for age-dependent changes in B cell homeostasis remain to be identified but will be important questions for future study given the profound compartment-wide influence they wield.

## Acknowledgements

The authors acknowledge financial support from the MRC (MR/P011225/1 to BS) and the National Institutes of Health (R01 AI093870 to AJY). We thank Fiona Powrie and the Oxford Centre for Microbiome Studies for germ-free mice.

## Author contributions

Conceived study; BS, AJY. Supervised experiments; BS. Supervised analyses; AJY. Performed experiments; MV. Provided germ-free mice; CP. Performed all mathematical modelling, coding, and statistical analyses; SR. Drafted manuscript; SR AJY BS. All authors approved the final version.

## Materials and Methods

### Model fitting, parameter estimation, and model selection

For pooled FM B cells, splenic GC B cells and lymph node GC B cells we fitted each mathematical model – schematically illustrated in Fig. 4A – simultaneously to the timecourses of total cell numbers, normalised chimerism, and Ki67 expression in host and donor cells. Mathematical and statistical methods are detailed in Text S3. Briefly, (i) we formulated the joint likelihood of the observations (Text S3.1), using the solutions of the model equations (Text S1), which in turn were functions of the model parameters and the empirical description of the timecourse of their putative precursor population (Text S4). (ii) We then used a Bayesian estimation approach with this likelihood and prior distributions of the model parameters to generate posterior distributions of these parameters (Text S3.2). Priors on the net loss rates of splenic and lymph node GC B cells were informed by data from the Ki67-YFP reporter mice (Text S6). (iii) The Bayesian approach also yielded a combined measure of the model’s quality of fit and its complexity (the Leave-One-Out Information Criterion, LOO-IC; Text S3.3). All code, data and prior distributions for parameters are available at https://github.com/sanketrane/B_cells_FM_GC. Model fits in Figs. 4, 5 and 6 were generated using the maximum *a posteriori* probability (MAP) estimates of the parameters, and are accompanied by envelopes that represent the spread of model predictions generated by sampling over these posterior distributions. Narrow envelopes therefore indicate that the model predictions are robust to variation in parameters; wide envelopes indicate sensitivity to parameter values.

### Mice

*Mki67*^*mCherry-CreERT2*^ mice were generated by targeted replacement of the terminal exon 14 of the *Mki67* locus with a modified exon 14 sequence with upstream FRT flanked neomycin cassette, and downstream mCherry fusion construct, IRES sequence and CreERT2 cDNA. Mice were crossed with *actin-FLPe* mice to excise the neomycin selection cassette, before crossing with *Rosa26*^*RYFP*^ strain (Srinivas et al., 2001), to generate *Mki67*^*mCherry-CreERT2*^ *Rosa26*^*RYFP*^ double reporter mice. Cre recombinase activity was induced *in vivo* in these mice following their i.p. injection with 2mg of tamxoxifen (Sigma) diluted in corn oil (Fisher Scientific) for five consecutive days.

Busulfan chimeras were generated as previously described (Hogan et al., 2015, 2017). C57Bl6/J mice were used as bone marrow donors and SJL.C57Bl6/J congenic mice as hosts. Donor bone marrow was obtained from femurs of age- and sex-matched C57Bl6/J mice. Thereafter, these bone marrow suspensions were depleted of T and B cells by immunomagnetic selection, using biotinylated antibodies to respectively CD3 (eBioscience, 1/500 dilution), TCR-beta (eBioscience, 1/500 dilution) and B220 (eBioscience, 1/200 dilution). Captured cells were bound to streptavidin-coupled Dynabeads (Life Technologies) and the unbound fraction depleted of mature T cells and B cells. 24 hours after the final busulfan injection, eight to ten million cells were injected i.v. in the busulfan-treated mice. At the indicated time points after BMT, host mice were sacrificed and spleen, lymph nodes and bone marrow were harvested and processed for further analysis. These mice, together with *Rag2*^*GFP*^ mice (Yu et al., 1999) and *Mki67*^*mCherry-CreERT2*^ *Rosa26*^*RYFP*^ double reporter mice were bred at Charles River U.K. Ltd and the Comparative Biology Unit, Royal Free Hospital. Experiments were performed according to the UCL Animal Welfare and Ethical Review Body and Home Office regulations. Germ-free mice were housed at the Oxford Centre for Microbiome Studies, Oxford, UK.

### Flow cytometry

Flow cytometric analyses were performed on 2 × 10^6^ cells from organs of interest. Cells were stained for 1 hour in the dark at 4°C with monoclonal antibodies (Abs) at a saturating concentration in 100 *µ*l of PBS. The following surface antigens were detected by the indicated mAb clone: B220 (RA3-6B2-BV785 and RA3-6B2-BV421, BioLegend), CD21 (7EG-PerCP-Cy5.5, Biolegend), CD23 (B3B4-FITC, Biolegend; B3B4-BUV737, BD Biosciences), CD45.1 (A20-BV650, Biolegend), CD45.2 (104-FITC, eBioscience; 104-PE-TR, Biolegend), CD93 (AA4.1-APC, Biolegend), CD95 (Jo2-biotin, BD Biosciences), GL7 (Ly77-PerCP-Cy5.5, Biolegend), IgD (11-26x.2a-BV421, Biolegend), IgM (Il/41-PE-Cy7, eBioscience) and live/dead Near-IR (Life Technologies). A secondary staining step was performed using streptavidin-BUV395 (BD Biosciences, 0.5 *µ*g/ml) or streptavidin-PerCP-Cy5.5 (BioLegend, 0.4 *µ*g/ml). Cells were stained for 30 min in the dark at 4°C. Subsequently, cells were washed in handling media, and immediately analysed by flow cytometry. For intracellular staining, cells were fixed and permeabilised using the FoxP3/transcription factor staining buffer set (eBioscience). Ki67 was detected using SolA15-FITC or SolA15-PE (eBioscience). Unless otherwise stated, individual populations were electronically gated as; splenic T1 cells, B220^hi^ AA4.1^pos^ IgM^hi^ CD23^low^; splenic and lymph node T2/3 cells, B220^hi^ AA4.1^pos^ CD23^hi^; FM B cells, B220^hi^ AA4.1^neg^ CD23^hi^; and GC B cells, B220^hi^ GL7^hi^. Data were analysed using FlowJo v10 (Becton Dickinson & Company).

## Supporting Information

**Fig. S1.**
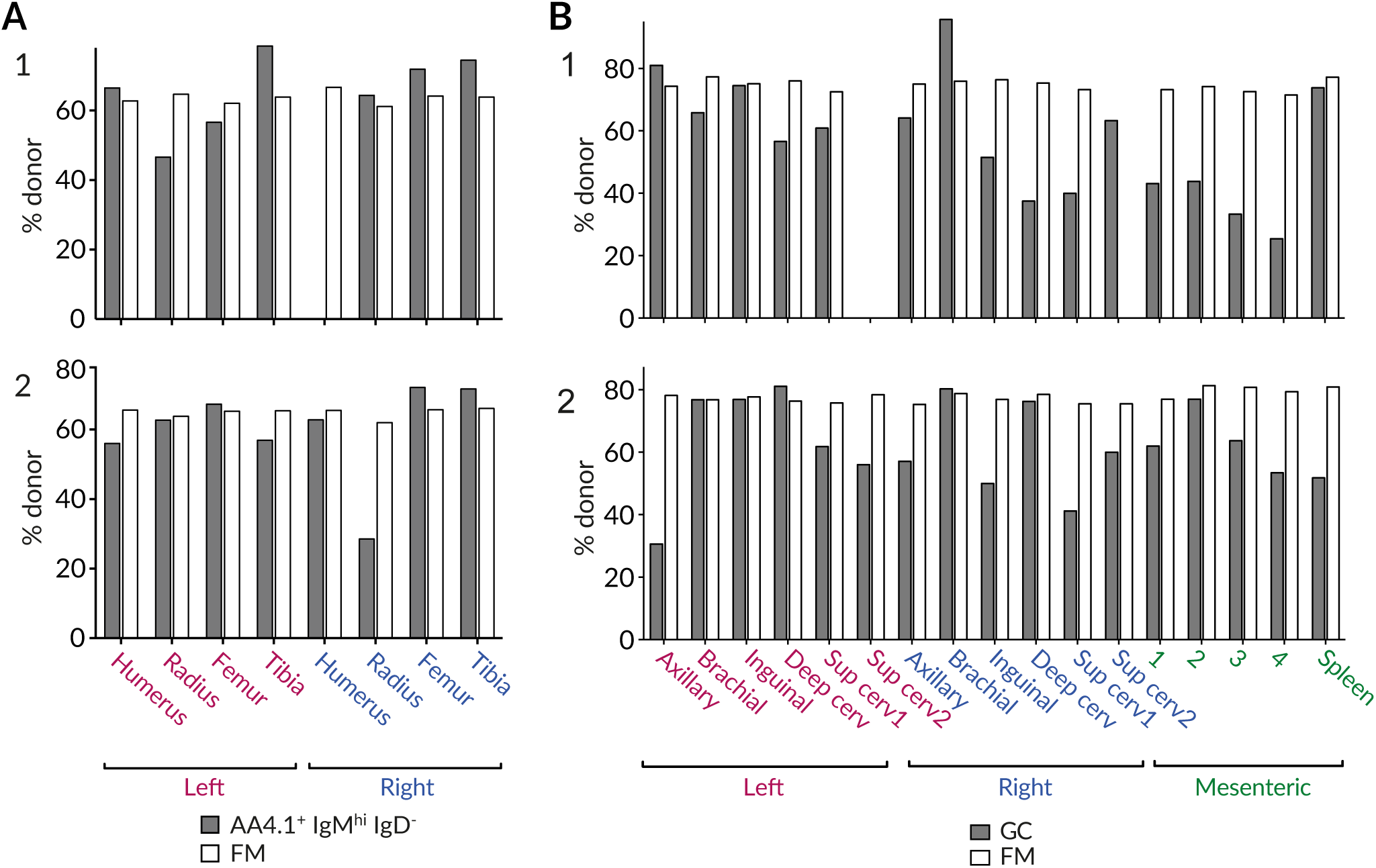
Variation in the degree of stable chimerism in B cell subsets within the same mice. **(A)** Variation in the chimerism among AA4.1^+^IgM^hi^IgD^−^ B cell progenitors and recirculating FM B cells within different BM sites. **(B)** Variation in chimerism across different lymph nodes. Data from two representative animals.

**Fig. S2.**
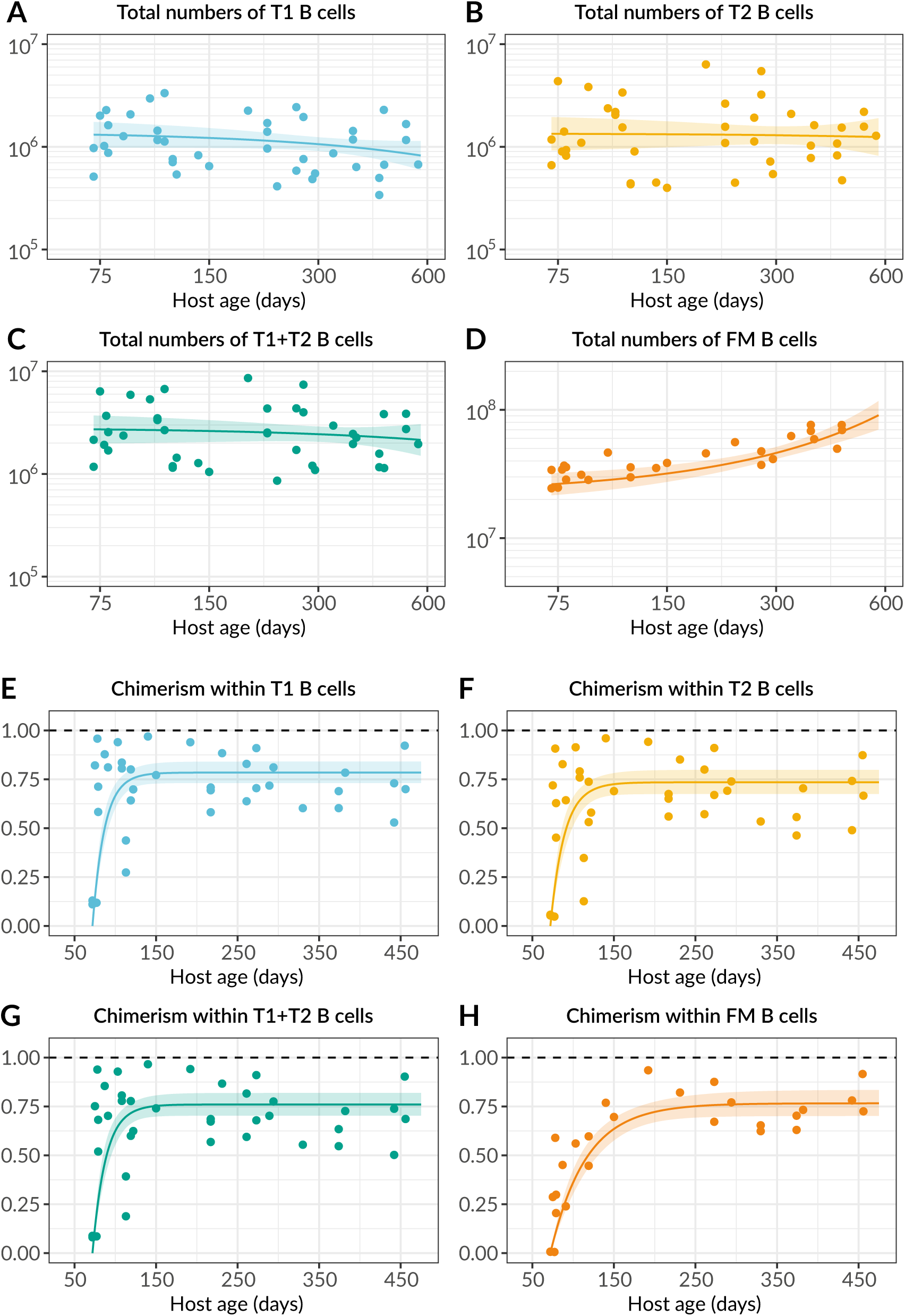
Empirical descriptions of the timecourses of total counts (A-D) and chimerism (E-H) in potential precursor populations.

**Fig. S3.**
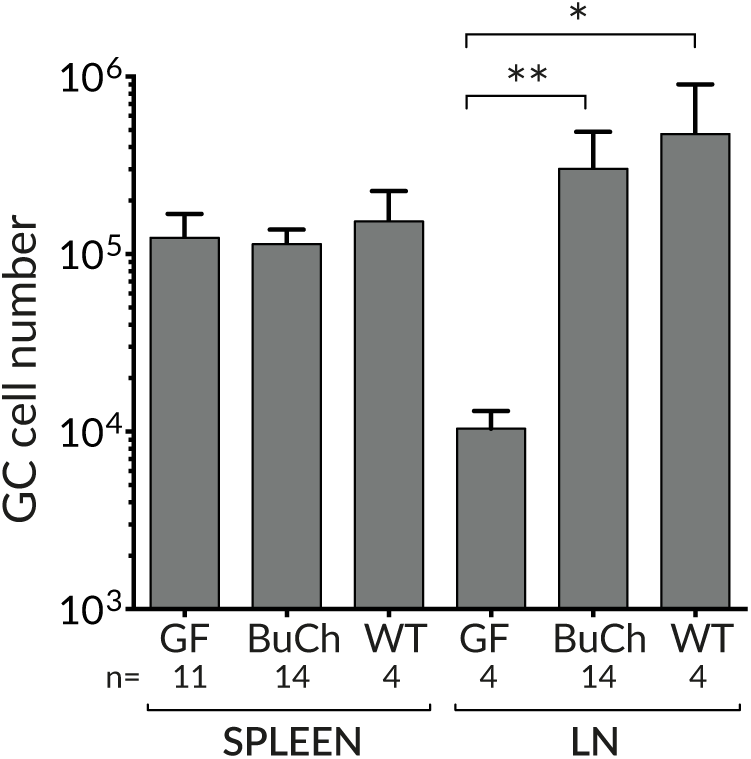
Comparing germinal centre B cell numbers in germ free, busulfan chimeric and WT mice. Bar chart shows absolute numbers of GC B cells in spleen and total LN from germ free (GF), busulfan chimeras (BuCh), and WT controls from a conventional barrier facility. Groups were of ages 10 weeks, 10-20 weeks and 10-14 weeks respectively. * p<0.05, ** p<0.01.

**Fig. S4.**
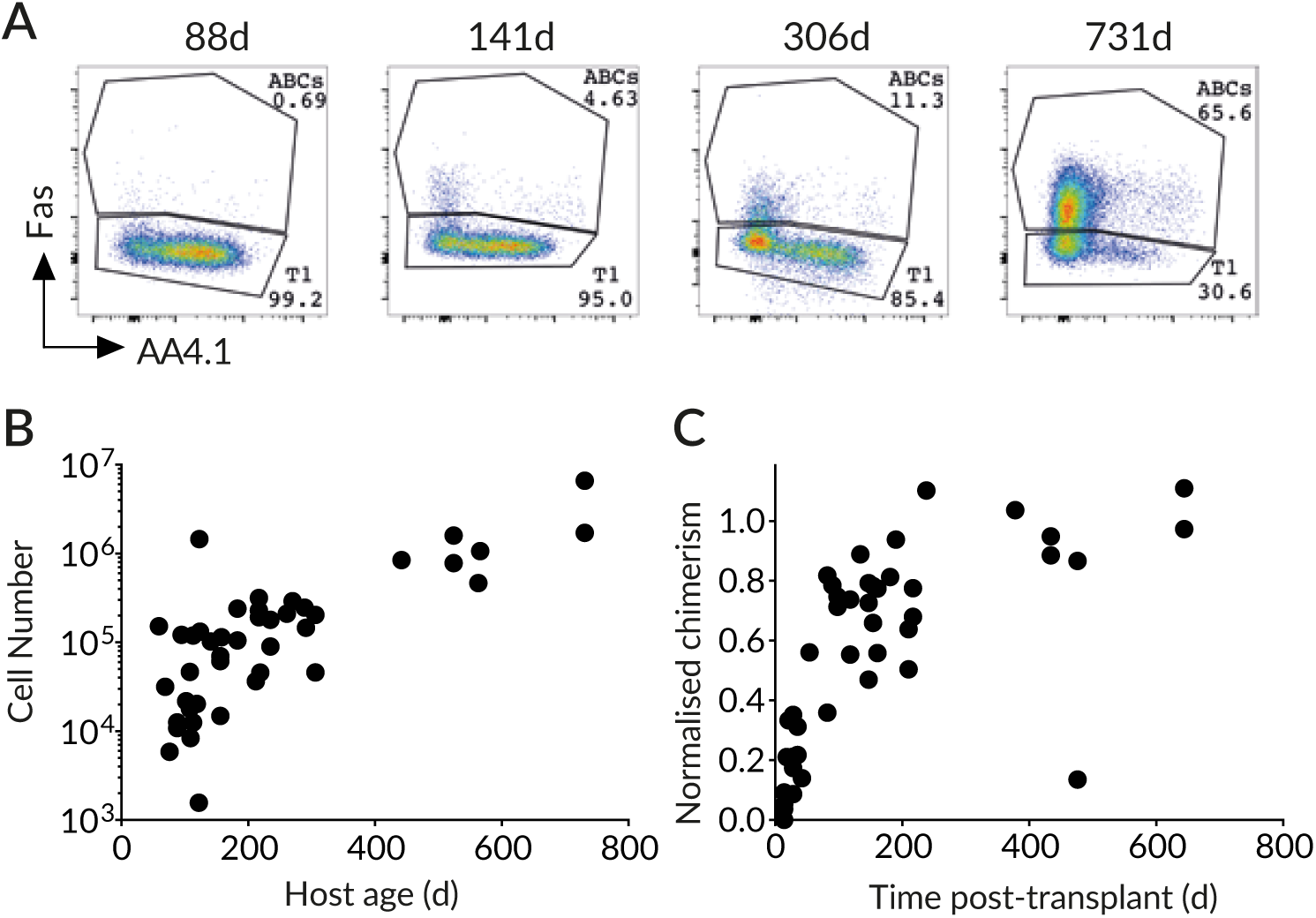
Age-associated B cells (AABC) accumulate in older mice. **(A)** Gating strategy for AABC. **(B)** AABC increase in number with host age. **(C)** AABC turn over within ∼300 days.

**Table S1.** Comparison of models describing the population dynamics of Follicular Mature (FM) B cells, pooled from LN and spleen, using Akaike weights (Text S3) as percentage measures of relative support. The strongly favoured model is highlighted.

| Source | Model and Akaike weight (%) |  |  |  |  |
| --- | --- | --- | --- | --- | --- |
|  | Simple homogeneous | Time-dependent turnover | Time-dependent division | Time-dependent influx | Kinetic heterogeneity |
| T1 | 9.0 | 64 | 3.0 | 18 | 5.0 |
| T2 | 0.0 | 0.0 | 0.0 | 0.0 | 0.0 |
| T1 + T2 | 0.0 | 0.5 | 0.0 | 0.5 | 0.0 |

**Table S2.** Comparison of models describing the population dynamics of germinal centre (GC) B cells in spleen, with or without the additional information regarding proliferation from the Ki67 reporter mice. ^*^Showed weak time-dependence in the rates of turnover (halving every ∼800 mo) or division (doubling every >1000 mo); were rejected in favour of the simple homogeneous model.

| Source | Model and Akaike weight (%) |  |  |  |  |  |
| --- | --- | --- | --- | --- | --- | --- |
|  | Simple homogeneous | Time-dependent turnover | Time-dependent division | Time-dependent influx | Kinetic heterogeneity | Incumbent |
| Without informing models using data from Ki67-Cre-ER<sup>T2</sup>-YFP reporter mice |  |  |  |  |  |  |
| T1 | 18 | 16 | 7 | 12 | 5 | 18 |
| T2 | 5 | 11 | 1 | 2 | 1 | 4 |
| Informing models using data from Ki67-Cre-ER<sup>T2</sup>-YFP reporter mice |  |  |  |  |  |  |
| T1 | 0 | 3 | 2 | 9 | 0 | 0 |
| T2 | 2 | 16* | 11* | 57 | 0 | 0 |

**Table S3.** Comparison of models describing the population dynamics of GC B cells in lymph nodes.

| Source | Model and Akaike weight (%) |  |  |  |  |  |
| --- | --- | --- | --- | --- | --- | --- |
|  | Simple homogeneous | Time-dependent influx | Time-dependent turnover | Time-dependent division | Kinetic heterogeneity | Incumbent |
| Without informing models using data from Ki67-Cre-ER<sup>T2</sup>-YFP reporter mice |  |  |  |  |  |  |
| T1 | 0 | 21 | 1 | 0 | 0 | 0 |
| T2 | 0 | 22 | 0 | 0 | 0 | 0 |
| FM | 6 | 37 | 11 | 2 | 0 | 0 |
| Informing models using data from Ki67-Cre-ER<sup>T2</sup>-YFP reporter mice |  |  |  |  |  |  |
| T1 | 0 | 0 | 0 | 0 | 5 | 0 |
| T2 | 0 | 0 | 0 | 0 | 11 | 0 |
| FM | 0 | 0 | 0 | 0 | 84 | 0 |

## Text S1 Mathematical models

### Simple homogeneous model

In this model we assume that cells form a kinetically homogeneous population that self-renews through homeostatic division with first-order kinetics at rate *α*, and is lost (turns over) at a rate *δ*, which combines death and onward differentiation. The inverse of *α* is the mean interdivision time, and the inverse of *δ* is the mean residence time of a cell. Influx of cells from the source compartment is denoted *ϕ*(*t*), which is the product of the *per capita* rate of influx *ψ* and the timecourse of the size of precursor population *S*(*t*), which is described empirically (see Text S4). We model the dynamics of Ki67^hi^ (*H*) and Ki67^lo^ (*L*) cells using the following ODE model;

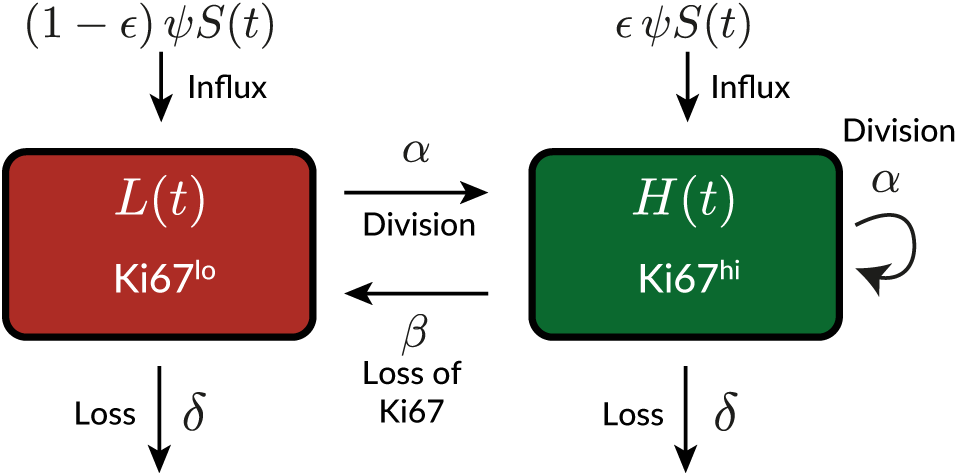

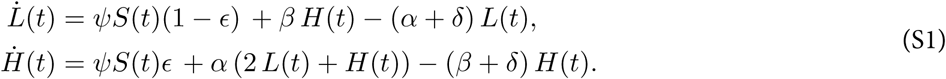

Here, *β* is the rate of loss of Ki67 expression after mitosis, and *ϵ* is the proportion of the cells entering from the source that are Ki67^hi^; we used its average observed value in the model fitting process. We assumed eqns. S1 held identically for host and donor cells. We fitted the following combinations of the solutions to these equations simultaneously to the timecourses of

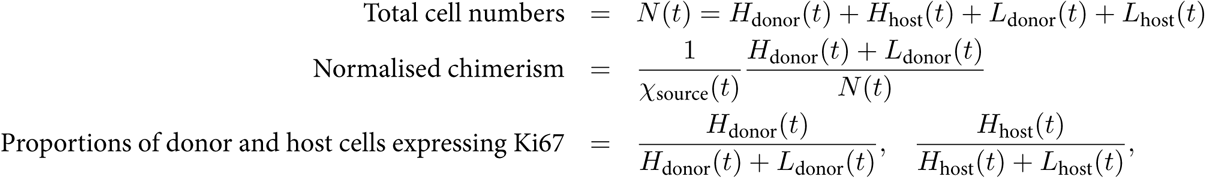

using the empirical descriptions of the size (*S*(*t*)) and chimerism (*χ*_source_(*t*)) of the source population (see Text S4). We define time *t*_0_ to be age at BMT of the youngest recipient (approximately 7 weeks), at which time the size of donor compartment is zero. Therefore, the Ki67^hi^ proportion among donor cells at *t*_0_ reflects that in the source, *κ*_donor_(*t*_0_) = *ϵ*. The Ki67^hi^ proportion among host cells at *t*_0_ is defined as *κ*_0_. We estimated *β, α, δ, ψ, κ*_0_ and *H*(*t*_0_) + *L*(*t*_0_), the size of the host compartment at *t*_0_.

### Time-dependent models

In these extensions of the model above, the *per capita* rate of influx of new cells from the source population *ψ*, the rate of cell division *α*, or the rate of loss *δ* may vary with time. We allowed each sub-model to exhibit time-dependence in only one process. These three sub-models are also homogeneous; at any given instant, all cells in the population exhibit the same rates of division and turnover. We assumed that the time-dependent parameter varied with mouse age *t* as exp(*rt*), where *r* was estimated from the data and was unconstrained (*i.e.* the rate constant could either rise or fall with time).

### Kinetic-heterogeneity model

This model comprises two subsets, which are independent, fed separately from the same source population, and are lost and/or divide at different rates. The resulting dynamics of the population as a whole are therefore the weighted average of the more ‘transient’ subset (rapid net loss, *δ*_*f*_ − *α*_*f*_) and a ‘persistent’ subset (slower net loss, *δ*_*s*_ − *α*_*s*_). We solve the following equations for Ki67^hi^ and Ki67^lo^ cells among the transient and persistent subsets, and formulate it identically for host and donor cells;

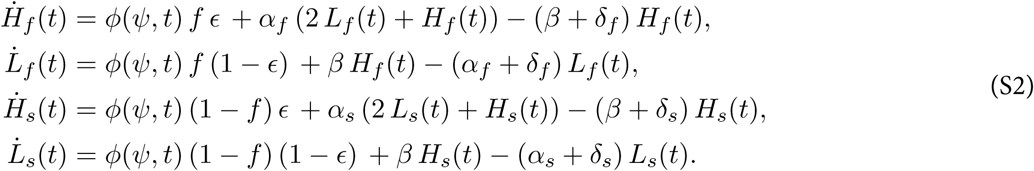

Along with the kinetic parameters we also estimate the proportions of the transient subset in the precursor (*f*) and in the target (*q*) populations, and the initial fractions of Ki67^hi^ cells in the transient and persistent subsets. The initial numbers of host-derived cells in the transient and persistent subsets are defined as *N*_0_ *q* and *N*_0_ (1 − *q*), respectively.

### Incumbent model

In this model, described in Hogan et al. (2015) and Rane et al. (2018), heterogeneity is exhibited only in the host compartment, which is assumed to comprise (i) an ‘incumbent’ subset of older, self-renewing cells that are resistant to displacement by new cells and (ii) a ‘displaceable’ subset that is replaced continuously by cohorts of new cells entering the pool. All donor cells are assumed to behave as displaceable cells.

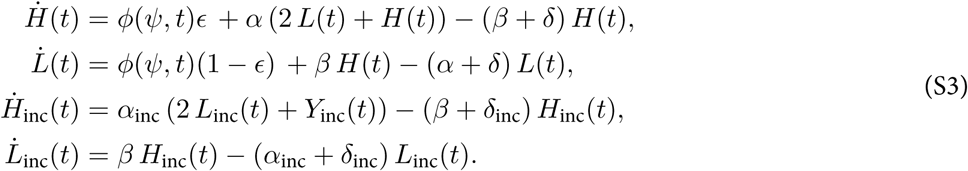

We assume that the incumbent subset is established early in life, before the minimum age of BMT in our chimeric animals (∼7 weeks).

## Text S2 The time taken to approach to stable chimerism in a B cell population is determined predominantly by the clonal lifetime

Here we illustrate for the simplest homogenous model the factors that determine the rate at which chimerism in a population reaches that of its precursor population. Assume a population *N* (*t*) is fed by precursors at constant total rate *ϕ*, divides at *per capita* rate *α* and is lost through death or differentiation at *per capita* rate *δ*;

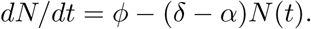

The quantity *δ* − *α* is the net loss rate, which we denote λ:

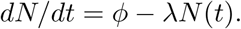

Assume the source acquires a stable chimerism *χ*, and that host (*h*) and donor (*d*) cells behave identically;

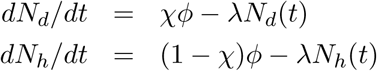

The normalised chimerism of the population is

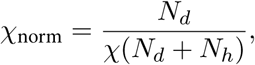

which evolves according to

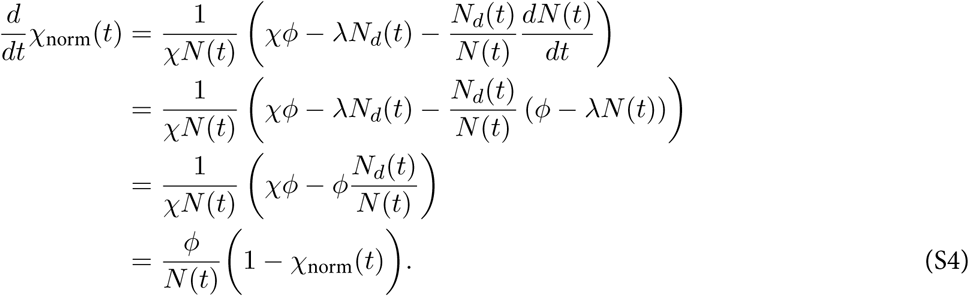

If the population is at equilibrium then *N* (*t*) = *ϕ/*λ, giving

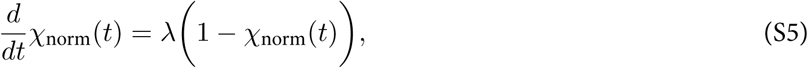

which implies

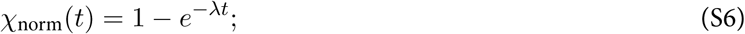

that is, the chimerism in the population reaches that of its precursors at a rate determined purely by the clonal lifespan 1*/*λ. If the population is initially out of equilibrium at size *N*_0_,

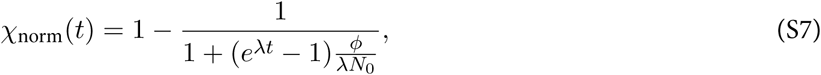

the rate of approach to *χ*_norm_ = 1 is then governed by both λ and the daily influx as a proportion of the initial pool size, *ϕ/N*_0_ (intuitively, if the pool is initially over-populated, *ϕ/*λ*N*_0_ < 1 and chimerism increases more slowly because of the excess of host cells; if the pool is depleted, *ϕ/*λ*N*_0_ > 1 and stable chimerism is achieved more quickly). Equation S7 reduces to S6 when *N*_0_ = *ϕ/*λ.

## Text S3 Fitting and selecting mathematical models

### S3.1 Likelihood

We attempted to explain the kinetics of host and donor cells in busulfan chimeric mice with an array of mathematical models, detailed in the main text and illustrated in Fig. 4A. As described, variation in the degree of depletion of host HSCs by busulfan treatment led to mouse-to-mouse variation in the level of stable bone-marrow chimerism (the fraction that are donor-derived), and hence also in peripheral subsets. We removed this variation by dividing the chimerism in each B cell subset by the chimerism *χ* in the T1 precursor population. This normalised chimerism (donor fraction) is

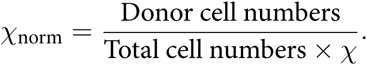

This approach allows us to fit a single model to data from multiple mice. Each model was fitted simultaneously to the timecourses of total cell counts (*N* (*t*), the sum of host and donor cells), the normalised chimerism *χ*_norm_(*t*), and the proportions of Ki67^hi^cells in the host and donor compartments (*κ*_host_(*t*) and *κ*_donor_(*t*)). Cell counts were log-transformed while *χ*_norm_, *κ*_host_ and *κ*_donor_ were logit-transformed, to ensure that measurement errors were approximately normally distributed. The joint likelihood of the datasets (with variables representing their transformed values) is then

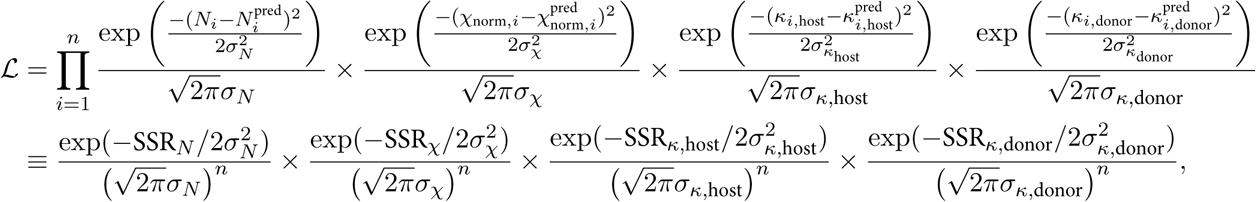

where *n* is the number of animals, each yielding four observations, and SSR denotes the sum of squared residuals, with each being a function of the data and the model parameters. This gives the joint log-likelihood (up to a constant);

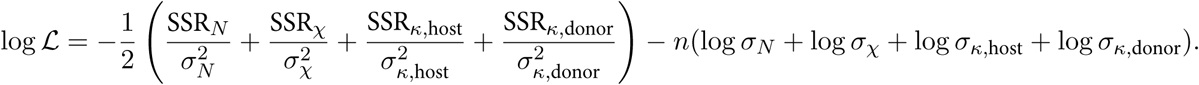

### S3.2 Parameter estimation

We used a Bayesian approach to estimating the model parameters, the errors associated with the measurements in each dataset, and a measure of support for each model. The inputs to this procedure are the joint likelihood shown above, and a set of prior distributions on the model parameters and the unknown measurement errors in each dataset. We refer to these unknowns collectively as *θ*. The Bayesian procedure updates these priors with the likelihood, to generate posterior distributions of *θ* that reflect our knowledge of these parameters in the light of the data, collectively denoted *y*. Strong (narrow) priors help to regularise a model’s behaviour and prevent it from learning too much from the data – and hence guard against over-fitting. The joint posterior distribution of the parameters is calculated using Bayes’ rule,

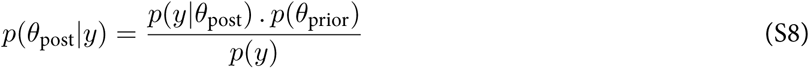

where *p*(*y*) is the likelihood of the data (averaged over the priors) that normalises the posterior such that it integrates to 1. We consider priors to be tools that improve a model’s ability to learn from the data, and subjected them to similar standards of evaluation and revaluation as any other component of the model. Detailed descriptions of the priors, together with the code and data for performing all of the analyses presented in this study, are available at https://github.com/sanketrane/B_cells_FM_GC.

The models were represented as systems of ordinary differential equations (ODEs), described in detail in Text S1. We solved them numerically using the *integrate_ode_rk45* solver in the *Stan* programming language and used the default no-U-turn sampler (NUTS) to generate the posterior distributions of the parameters. We confirmed that the log-transformed cell counts, and the logit-transformed values of the normalised chimerism and the Ki67^hi^ proportions in host and donor compartments, were all normally distributed with constant errors (standard deviations). These standard deviations were additional parameters that were estimated from the data. We used the *R-stan* package in *R* to interface and compile the *Stan* scripts that encoded the priors, model definitions, sampling and fitting procedures.

### S3.3 Comparing models

The assessment of a model’s utility depends on how accurately it explains a given dataset (measured by the likelihood) and also how well it can predict new observations. A complex model with an excessive number of parameters will tend to overfit any given dataset and perform poorly when predicting new observations. On the other hand, a model that is too simple will fail to capture trends in the data, generate a low likelihood, and will also make poor predictions of new observations. The Akaike Information Criterion (Akaike, 1974, Burnham and Anderson, 2002) is commonly used to identify parsimonious models that trade off the likelihood and complexity. However, the AIC penalises all model parameters equally, which may not be appropriate when they differ in their ability to influence a fit. In this study, we use the Leave-one-out information criterion (LOO-IC; Vehtari et al. (2017)) which penalizes the addition of model parameters only to the extent that they contribute to overfitting.

Briefly, we define the log predictive density of a single observation *y*_*i*_ given a model with parameters *θ* – that is, the average value of the log-likelihood log (*p*(*y*_*i*_|*θ*)) across the joint posterior distribution of *θ*. We approximate this by making *D* draws from the posterior distribution, calculating the likelihood of *y*_*i*_ for each set of parameters, and averaging. This process is repeated for each data point (*y*_1_, …, *y*_*n*_) to calculate the log point-wise predictive density (lppd) for the whole dataset:

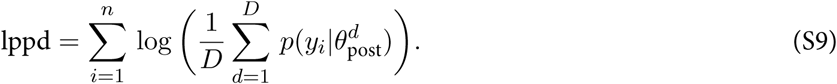

One then uses the leave-one-out (*loo*) method, a special case of cross-validation, whereby the dataset of *n* observations (*y*_1_, …, *y*_*n*_) is partitioned into *n* training datasets each of size *n* − 1. Fitting the model to the training sample that excludes datapoint *i* gives a joint posterior 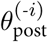. This posterior is then used to estimate the prediction accuracy of the model for the excluded observation *i* (the test sample), which is defined as the log of the average likelihood of the test sample across the posterior distribution. This likelihood is approximated by averaging over *D* draws from the posterior. This process is repeated, making each observation in the dataset (*y*_1_, …, *y*_*n*_) the test sample, and the lppd^*loo*^ is defined to be the sum of the log likelihoods of all these prediction accuracies:

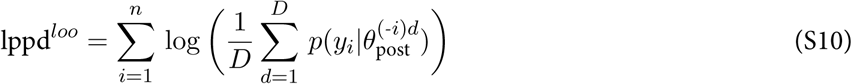

where the term in large parentheses characterizes the *D* posterior simulations fitted on *n* − 1 observations when the *i*^*th*^ observation is left out. The information criterion LOO-IC is defined as −2 × lppd (Vehtari et al., 2017). To calculate it we use the *loo-2.0* package in the *rstan* library, which estimates the lppd^*loo*^ using Pareto-smoothed importance sampling (PSIS) – an approximation of leave-one-out cross-validation that uses existing posterior draws from the model fits (Vehtari et al., 2015).

We then used the estimated LOO-IC values to assess the relative support for models using the analog of the Akaike weight – the probability that a given model will explain new data better than other models considered in the analysis. Following Burnham and Anderson (Burnham and Anderson, 2002), these weights are

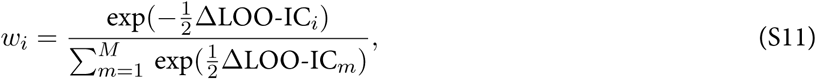

where ΔLOO-IC_*i*_ is the difference in LOO-IC values between model *i* of *M* candidates and the model with the lowest LOO-IC value.

## Text S4 Modelling the dynamics of precursor cell numbers and chimerism

We considered T1, T2, and T1+T2 combined as the potential direct precursors (sources) of FM B cells, and T1, T2 and FM B cells as potential sources of GC B cells. In adult mice, we described the time-variation in the sizes of these populations with the empirical descriptor function *S*(*t*) = *S*_0_ *e*^*−ν t*^ (Fig. S2, panels A-D), where the parameters *S*_0_ and *ν* were estimated by fitting to the log-transformed cell counts using least squares.

Similarly, the timecourses of donor chimerism in these populations were all described well with *χ*(*t*) = *χ*_stable_ (1 − *e*^*−ν t*^), shown in Fig. S2, panels E-H; here, *χ*_stable_ and *ν* were estimated using non-linear least squares.

We assumed a constant *per capita* rate of influx *ψ* from the source *S*(*t*), giving a total influx of *ϕ*(*t*) = *ψS*(*t*) cells/day. The daily influx of host and donor cells into the target population is then

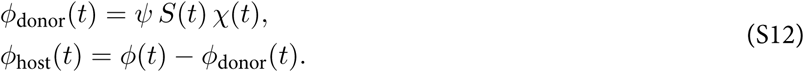

The unknown *ψ* is estimated along with the other model parameters. In the time-dependent recruitment model, we assumed the form *ψ*(*t*) = *ψ*_0_ *e*^*p t*^, and estimated *ψ*_0_ and *p*.

## Text S5 Modelling the development of the FM B cell pool in young mice

### S5.1 Empirical description of T1 precursor numbers in young mice

To capture the dynamics of T1 cells in young mice (Fig. 5A) we used the empirical function *S*(*t*) = *S*_0_ (1 + *t*^*n*^ exp(−*bt*)), and fitted this to the log-transformed cell counts, using least squares to estimate *S*_0_, *n* and *b*.

### S5.2 Explaining the developmental dynamics of FM B cells in young mice

To test the hypothesis of lower recruitment of T1 B cells in neonates than in adults, we allowed the rate of influx to increase with time early in life, approaching the value *ψ* estimated from our best-fitting model in adults aged 7 weeks and older; we assumed the form *ψ*(*t*) = *ψ*(1 − exp(−*r*_*ψ*_*t*)). The estimated rate *r*_*ψ*_ was sufficiently large that *ψ*(*t*) was very close to *ψ* at age 7 weeks (Fig. 5E).

To test whether FM B cells in young mice are lost more rapidly than those in adult mice, we extended the time-dependent loss model, in which we had described the loss rate from age *t*_0_=7 weeks onwards as 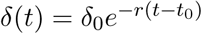. For *t* < *t*_0_ we assumed 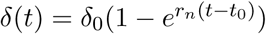 with *r*_*n*_ > *r* (Fig. 5F).

We fitted both extensions of the time-dependent loss model to the counts of FM B cells in young mice separately, estimating *r*_*ψ*_ and *r*_*n*_ in the process.

### S5.3 Estimating the age distribution of FM B cells in young mice

To generate the predicted age distributions of cells under the two models above, we recast them as partial differential equations (PDEs) that explicitly track cell age. In the time-dependent loss model, the population density of FM B cells of age *a* in mice of age *t* is given by the solution to

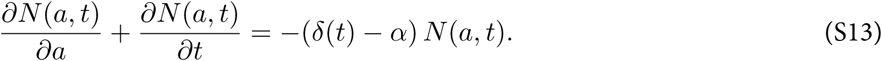

The rate of influx of cells of age zero *N* (0, *t*) is the source influx *ψS*(*t*), or *ψ*(*t*)*S*(*t*) for the model in which the *per capita* influx rate varies with age. The other boundary condition is the age distribution of cells at time zero, size *N* (*a*, 0). We assumed that the FM B cell compartment at the time of birth is sufficiently small that we could set *N* (*a*, 0) = 0. We solved this model using the parameters estimated from fitting the extensions of time-dependent loss model (described above) to the total counts of FM B cells in young mice. We then calculated the normalised cell age distribution of FM B cells at *t* = 7 weeks using

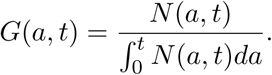

See Rane et al. (2018) for full details of the solution of this class of model.

## Text S6 Using information from Ki67-reporter mice to aid discrimination between models of GC B cell dynamics

The Ki67-Cre-ER^*T*2^-YFP system allows us to track cohorts of cells that underwent cell division during tamoxifen treatment. We measured the frequencies of YFP-expressing cells at day 4 and day 62 post-tamoxifen and, for each model (as described below), used the decline in YFP expression over this time period to constrain the rates of loss and/or division. YFP expression is preserved upon cell division but is diluted by loss or onward differentiation. To illustrate, for the simple homogeneous model, YFP expression will decline at the net rate of loss of the population λ, which is *δ* − *α*. We can therefore relate λ to the fold loss of YFP expression over a time *τ*:

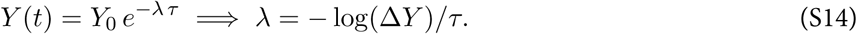

### Priors on Δ*Y*

We took the strategy of making Δ*Y* a parameter in the model, using its observed values to generate its prior; and sampling from this prior then allowed us to estimate or constrain other parameters. For splenic GC B cells, the mean YFP-labelled fraction dropped from 0.35 to 0.04 over 8 weeks, yielding Δ*Y* ∼ 0.12. This, together with the scatter in Δ*Y* observed in YFP reporter mice (4 mice at the 2 week timepoint and 5 mice at the 8 week timepoint, unpaired data; Fig. 6D in the text), suggested Δ*Y* ∼ 𝒩(0.12, 0.05). Lymph node GC B cells exhibited Δ*Y* ≃ 0.42. When assuming T1 or T2 as their precursors, which turn over rapidly and are therefore not expected to provide a persistent source of YFP-labelled cells after withdrawal of tamoxifen, we therefore assumed Δ*Y* ∼ 𝒩(0.42, 0.05). When assuming FM B cells to be precursors, which turn over more slowly, we considered the possibility that FM B cells might act as a reservoir that feeds new YFP^+^ cells into LNGC for some time after withdrawal of tamoxifen. In the case the drop in YFP expression yields only a lower bound on λ. Accordingly, we assumed that Δ*Y* was skew-normally distributed with a bias towards values less than the mean of 0.42, and a standard deviation of 0.1 (Δ*Y* ∼ SkewNormal(0.42, 0.1, −5)). Specifically, if *µ* ∈ ℝ, *σ* ∈ ℝ^+^, and *k* ∈ ℝ, then for y ∈ ℝ,

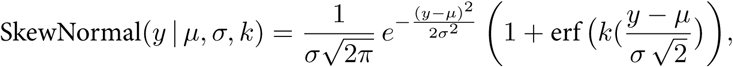

where ‘erf’ is the Gaussian error function. For each model we used the YFP information in the following ways:

### Simple homogeneous model (with or without time-dependent influx)

Using the above priors on Δ*Y* and the division rate *α*, we then estimated the rate of loss (*δ*) using equation S14;

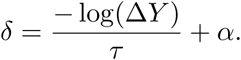

### Time-dependent division or loss

With time dependent division, we assumed priors for Δ*Y* and *δ* and calculated the rate of division at *t*_0_, which we denote *α*_0_. With time dependent loss loss, we assumed priors for Δ*Y* and *α* and calculated the rate of loss at *t*_0_ (*δ*_0_):

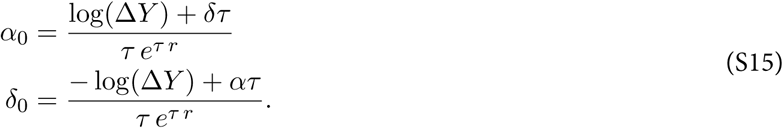

### Kinetic heterogeneity

This model (eqns. S2) predicts a biphasic loss of YFP, reflecting the net loss rates of the transient (λ_*f*_) and persistent (λ_*s*_) subsets and which were present at unknown frequencies *q* and 1 − *q*, respectively. We then assigned priors Δ*Y*, λ_*f*_ and λ_*s*_, to give *q*:

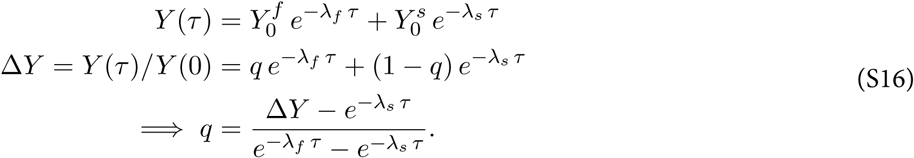

Here 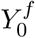 and 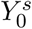 are the fractions of YFP-labelled cells in the transient and persistent subsets, respectively, at time *τ* = 0. Assuming λ_*f*_ > λ_*s*_, the constraint 0 < *q* < 1 in turn constrains the priors on λ_*f*_ and λ_*s*_;

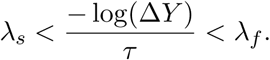

Placing priors on the division rates *α*_*f*_ and *α*_*s*_, we could then calculate the rates of loss of transient (*δ*_*f*_) and persistent subsets (*δ*_*s*_) using

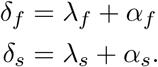

### Incumbent

We derived a similar relationship between λ_displaceable_ and λ_incumbent_ to that in equation S16,

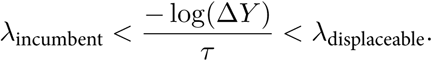

Using priors on Δ*Y*, λ_displaceable_, λ_incumbent_, *α*_inc_ and *α*, we calculated *δ* and *δ*_inc_:

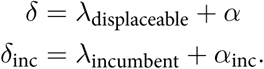

## References

Akaike, H. (1974). A new look at the statistical model identification. IEEE Transactions on Automatic Control 19, 716–723.

Allman, D., Lindsley, R. C., DeMuth, W., Rudd, K., Shinton, S. A. and Hardy, R. R. (2001). Resolution of three non-proliferative immature splenic B cell subsets reveals multiple selection points during peripheral B cell maturation. Journal of Immunology 167, 6834–6840.

Anderson, S. M., Khalil, A., Uduman, M., Hershberg, U., Louzoun, Y., Haberman, A. M., Kleinstein, S. H. and Shlomchik, M. J. (2009). Taking advantage: high-affinity B cells in the germinal center have lower death rates, but similar rates of division, compared to low-affinity cells. J Immunol 183, 7314–25.

Basso, K. and Dalla-Favera, R. (2015). Germinal centres and B cell lymphomagenesis. Nature Reviews Immunology 15, 172–184.

Bergqvist, P., Stensson, A., Hazanov, L., Holmberg, A., Mattsson, J., Mehr, R., Bemark, M. and Lycke, N. Y. (2013). Re-utilization of germinal centers in multiple Peyer’s patches results in highly synchronized, oligoclonal, and affinity-matured gut IgA responses. Mucosal Immunol 6, 122–35.

Burnham, K. P. and Anderson, D. R. (2002). Model Selection and Multimodel Inference: A Practical Information-Theoretic Approach. Second edition, Springer-Verlag, New York, USA.

Cariappa, A. (2009). The follicular versus marginal zone B lymphocyte cell fate decision. Nature Reviews Immunology 9, 767–777.

De Boer, R. J., Mohri, H., Ho, D. D. and Perelson, A. S. (2003). Estimating average cellular turnover from 5-bromo-2’-deoxyuridine (BrdU) measurements. Proc R Soc Lond B Biol Sci 270, 849–58.

De Boer, R. J. and Perelson, A. S. (2013). Quantifying T lymphocyte turnover. J Theor Biol 327, 45–87.

De Silva, N. S. and Klein, U. (2015). Dynamics of B cells in Germinal Centres. Nature Reviews Immunology 15, 137–148.

Figge, M. T. (2005). Stochastic discrete event simulation of germinal center reactions. Phys Rev E Stat Nonlin Soft Matter Phys 71, 051907.

Förster, I. and Rajewsky, K. (1990). The bulk of the peripheral B-cell pool in mice is stable and not rapidly renewed from the bone marrow. Proceedings of the National Academy of Sciences 87, 4781–4784.

Frasca, D., Diaz, A., Romero, M., Landin, A. M. and Blomberg, B. B. (2011). Age effects on B cells and humoral immunity in humans. Ageing Research Reviews 10, 330–335.

Fulcher, D. A. and Basten, A. (1997). Influences on the lifespan of B cell subpopulations defined by different phenotypes. European Journal of Immunology 27, 1188–1199.

Gibson, K. L., Wu, Y.-C., Barnett, Y., Duggan, O., Vaughan, R., Kondeatis, E., Nilsson, B.-O., Wikby, A., Kipling, D. and Dunn-Walters, D. K. (2009). B-cell diversity decreases in old age and is correlated with poor health status. Aging Cell 8, 18–25.

Gossel, G., Hogan, T., Cownden, D., Seddon, B. and Yates, A. J. (2017). Memory CD4 T cell subsets are kinetically heterogeneous and replenished from naive T cells at high levels. eLife 6, 596.

Grabstein, K. H., Waldschmidt, T. J., Finkelman, F. D., Hess, B. W., Alpert, A. R., Boiani, N. E., Namen, A. E. and Morrissey, P. J. (1993). Inhibition of murine B and T lymphopoiesis in vivo by an anti-interleukin 7 monoclonal antibody. Journal of Experimental Medicine 178, 257–264.

Hao, Z. and Rajewsky, K. (2001). Homeostasis of peripheral B cells in the absence of B cell influx from the bone marrow. Journal of Experimental Medicine 194, 1151–1164.

Hogan, T., Gossel, G., Yates, A. J. and Seddon, B. (2015). Temporal fate mapping reveals age-linked heterogeneity in naive T lymphocytes in mice. Proceedings of the National Academy of Sciences 112, E6917–E6926.

Hogan, T., Nowicka, M., Cownden, D., Pearson, C., Yates, A. J. and Seddon, B. (2019). Differential impact of self and environmental antigens on the ontogeny and maintenance of CD4+ T cell memory. eLife 10.7554/eLife.48.

Hogan, T., Shuvaev, A., Commenges, D., Yates, A., Callard, R., Thiébaut, R. and Seddon, B. (2013). Clonally diverse T cell homeostasis is maintained by a common program of cell-cycle control. Journal of Immunology 190, 3985–3993.

Hogan, T., Yates, A. and Seddon, B. (2017). Generation of busulfan chimeric mice for the analysis of T cell population dynamics. Bio-Protocol 7, 1–8.

Kepler, T. B. and Perelson, A. S. (1993). Cyclic re-entry of germinal center B cells and the efficiency of affinity maturation. Immunol Today 14, 412–5.

Kline, G. H., Hayden, T. A. and Klinman, N. R. (1999). B cell maintenance in aged mice reflects both increased B cell longevity and decreased B cell generation. J Immunol 162, 3342–9.

Lam, K. P., Kühn, R. and Rajewsky, K. (1997). In vivo ablation of surface immunoglobulin on mature B cells by inducible gene targeting results in rapid cell death. Cell 90, 1073–1083.

Loder, F., Mutschler, B., Ray, R. J., Paige, C. J., Sideras, P., Torres, R., Lamers, M. C. and Carsetti, R. (1999). B cell development in the spleen takes place in discrete steps and is determined by the quality of B cell receptor-derived signals. Journal of Experimental Medicine 190, 75–89.

Martin, B., Bécourt, C., Bienvenu, B. and Lucas, B. (2006). Self-recognition is crucial for maintaining the peripheral CD4+ T-cell pool in a nonlymphopenic environment. Blood 108, 270–7.

Mesin, L., Ersching, J. and Victora, G. D. (2016). Germinal Center B Cell Dynamics. Immunity 45, 471–482.

Meyer-Bahlburg, A., Andrews, S. F., Yu, K. O. A., Porcelli, S. A. and Rawlings, D. J. (2008). Characterization of a late transitional B cell population highly sensitive to BAFF-mediated homeostatic proliferation. Journal of Experimental Medicine 205, 155–168.

Meyer-Hermann, M., Mohr, E., Pelletier, N., Zhang, Y., Victora, G. D. and Toellner, K.-M. (2012). A theory of germinal center B cell selection, division, and exit. Cell Rep 2, 162–74.

Miller, I., Min, M., Yang, C., Tian, C., Gookin, S., Carter, D. and Spencer, S. L. (2018). Ki67 is a graded rather than a binary marker of proliferation versus quiescence. Cell Reports 24, 1105–1112.e5.

Naradikian, M. S., Hao, Y. and Cancro, M. P. (2016). Age-associated B cells: key mediators of both protective and autoreactive humoral responses. Immunol Rev 269, 118–129.

Petro, J. B., Gerstein, R. M., Lowe, J., Carter, R. S., Shinners, N. and Khan, W. N. (2002). Transitional type 1 and 2 B lymphocyte subsets are differentially responsive to antigen receptor signaling. J Biol Chem 277, 48009–48019.

Rane, S., Hogan, T., Seddon, B. and Yates, A. J. (2018). Age is not just a number: Naive T cells increase their ability to persist in the circulation over time. PLoS Biology 16, e2003949–20.

Reboldi, A. and Cyster, J. G. (2016). Peyer’s patches: organizing B-cell responses at the intestinal frontier. Immunol Rev 271, 230–245.

Robert, P. A., Rastogi, A., Binder, S. C. and Meyer-Hermann, M. (2017). How to Simulate a Germinal Center. Methods Mol Biol 1623, 303–334.

Schwickert, T. A., Lindquist, R. L., Shakhar, G., Livshits, G., Skokos, D., Kosco-Vilbois, M. H., Dustin, M. L. and Nussenzweig, M. C. (2007). In vivo imaging of germinal centres reveals a dynamic open structure. Nature 446, 83–7.

Shahaf, G., Johnson, K. and Mehr, R. (2006). B cell development in aging mice: lessons from mathematical modeling. Int Immunol 18, 31–9.

Shulman, Z., Gitlin, A. D., Targ, S., Jankovic, M., Pasqual, G., Nussenzweig, M. C. and Victora, G. D. (2013). T follicular helper cell dynamics in germinal centers. Science 341, 673–7.

Srinivas, S., Watanabe, T., Lin, C. S., William, C. M., Tanabe, Y., Jessell, T. M. and Costantini, F. (2001). Cre reporter strains produced by targeted insertion of EYFP and ECFP into the ROSA26 locus. BMC Dev Biol 1, 4.

Srivastava, B., Quinn, W. J., Hazard, K., Erikson, J. and Allman, D. (2005). Characterization of marginal zone B cell precursors. Journal of Experimental Medicine 202, 1225–1234.

Su, T. T. and Rawlings, D. J. (2002). Transitional B lymphocyte subsets operate as distinct checkpoints in murine splenic B cell development. Journal of Immunology 168, 2101–2110.

Torres, R. M., Flaswinkel, H., Reth, M. and Rajewsky, K. (1996). Aberrant B cell development and immune response in mice with a compromised BCR complex. Science 272, 1804–1808.

Vehtari, A., Gelman, A. and Gabry, J. (2015). Efficient implementation of leave-one-out cross-validation and WAIC for evaluating fitted Bayesian models. Statistics and Computing, 1507.04544 27, 1413–1432.

Vehtari, A., Gelman, A. and Gabry, J. (2017). Practical Bayesian model evaluation using leave-one-out cross-validation and WAIC. Stat Comput 27, 1413–1432.

Victora, G. D. and Mesin, L. (2014). Clonal and cellular dynamics in germinal centers. Current Opinion in Immunology 28, 90–6.

Wittenbrink, N., Klein, A., Weiser, A. A., Schuchhardt, J. and Or-Guil, M. (2011). Is there a typical germinal center? A large-scale immunohistological study on the cellular composition of germinal centers during the hapten-carrier-driven primary immune response in mice. J Immunol 187, 6185–96.

Yu, W., Nagaoka, H., Jankovic, M., Misulovin, Z., Suh, H., Rolink, A., Melchers, F., Meffre, E. and Nussenzweig, M. C. (1999). Continued RAG expression in late stages of B cell development and no apparent re-induction after immunization. Nature 400, 682–687.

